# Inference of pairwise interactions from strain frequency data across settings and context-dependent mutual invasibilities

**DOI:** 10.1101/2024.09.06.611626

**Authors:** Thi Minh Thao Le, Sten Madec, Erida Gjini

## Abstract

We propose a method to estimate pairwise strain interactions from population-level frequencies across different endemic settings. We apply the framework of replicator dynamics, derived from a multi-strain SIS model with co-colonization, to extract from 5 datasets the fundamental backbone of strain interactions. In our replicator, each pairwise invasion fitness explicitly arises from local environmental context and trait variations between strains. We adopt the simplest formulation for multi-strain coexistence, where context is encoded in basic reproduction number *R*_0_ and mean global susceptibility to co-colonization *k*, and trait variations *α*_*ij*_ capture pairwise deviations from *k*. We integrate *Streptococcus pneumoniae* serotype frequencies and serotype identities collected from 5 environments: epidemiological surveys in Denmark, Nepal, Iran, Brazil and Mozambique, and mechanistically link their distributions. Our results have twofold implications. First, we offer a new *proof-of-concept* in the inference of multi-species interactions based on cross-sectional data. We also discuss 2 key aspects of the method: the site ordering for sequential fitting, and stability constraints on the dynamics. Secondly, we effectively estimate at high-resolution more than 70% of the 92 × 92 pneumococcus serotype interaction matrix in co-colonization, allowing for further projections and hypotheses testing. We show that in these bacteria both within- and between- serotype interaction coefficients’ distribution emerge to be unimodal, their difference in mean broadly reflecting stability assumptions on serotype coexistence. This framework enables further model calibration to global data: cross-sectional across sites, or longitudinal in one site over time, - and should allow a more robust and integrated investigation of intervention effects in such biodiverse ecosystems.

## 1 Introduction

Diversity is ubiquitous in nature. Quantifying diversity, its generative processes and consequences is a major aim in biology today, with applications in ecology, conservation, epidemiology and biomedicine. Although our ability to gather diversity data has increased in recent years to unprecedented levels of resolution, our ability to interpret such data, and mechanistically explain and cross-link species abundances in different studies or systems remains limited. It has been long recognized that the structure and magnitude of species interactions can drastically affect the complexity and stability of ecosystems [May, 1972, McCann, 2000], and patterns of species abundances [Motomura, 1932, MacArthur, 1960, Cody and Diamond, 1975, Hubbell, 2011, Tokita, 2004, 2006, Yoshino et al., 2008, Jacquet et al., 2016]. However, uncovering the rules of interactions and of multispecies coexistence from data remains still challenging empirically.

One such example in the microbiology literature are the microbial species (OTU taxa) populating the human or murine gut. These are often tackled with generalized Lotka-Volterra systems applied to longitudinal time-series data of species abundances in the same host over time to infer pairwise species interactions [Stein et al., 2013, Marino et al., 2014, Coyte et al., 2015, Alshawaqfeh et al., 2017, Lo and Marculescu, 2017, Hu et al., 2022].

Another example, from the epidemiology literature, comes from multi-strain dynamics models, where epidemiological diversity and processes included in structural assumptions are used to describe interactions between strains and predict their population-level effects. Interactions can be included in the form of cross-immunity coefficients e.g in influenza [Koelle et al., 2006], pairwise scaling coefficients between strains in co-colonization vulnerabilities, e.g. in the context of pneumococcus serotypes [Gjini et al., 2016], and other life-history traits, inspiring also simulation-based inference studies in SIS endemic systems [Man et al., 2023, 2019]. Some other epidemiological models focus on interactions between two different diseases or pathogens, e.g. between influenza and pneumococcus [Shrestha et al., 2013].

More often than mechanistic models, statistical (non-mechanistic) models are used to infer strain interactions, based on Markov chain description of colonization dynamics in a cohort of individuals tracked over time to estimate conditional hazards for acquisition of different strain pairs [Lipsitch et al., 2012] or statistical odds-ratios, e.g. among Dengue serotypes [Aguas et al., 2019].

In general, so far, most existing methods for detecting multi-strain pathogen interactions have been limited to assuming uniform interactions across strains (i.e. all modeled strains interact with one another with the same coefficient), or described only marginal interactions between pairs of pathogen strains, without accounting for other co-occurring strains) [Alizon et al., 2019, Chaturvedi et al., 2011, Mejlhede et al., 2010, Auranen et al., 2010, Gjini, 2017].

Thus, numerical quantification of direct pairwise strain-interactions from *real* epidemiological data for a *high* strain number remains a hard and mostly untackled problem in the field of endemic microbial ecosystems. In this study, inspired by pneumococcus, we will propose a method for quantifying a high-dimensional interaction matrix between strains, from crosssectional relative abundance data, based on the replicator equation [Madec and Gjini, 2020, Hofbauer and Sigmund, 1998]. Our study contributes towards a deeper qualitative and quantitative understanding of the high-dimensional interaction landscape in multi-strain systems.

### Pneumococcus: a case study for linking different cross-sectional diversity data

A very well-studied interacting multi-strain system in epidemiology is the system of *Streptococcus pneumoniae* bacteria, their serotype structure, and their global epidemiology of colonization, and co-colonization. *Streptococcus pneumoniae* bacteria are typically asymptomatic colonizers of the human nasopharynx. Yet, it is known that carriage is a pre-requisite for disease, and that these bacteria can cause pneumonia, invasive disease, otitis, meningitis and other complications, in children or vulnerable adults. Single and multiple carriage and invasive pneumococcal disease are typically most prevalent in children under 5 years old, and vary across settings, depending on demographic and environmental factors, many of which are not understood [Dekaj and Gjini, 2024, Garcia Quesada et al., 2021].

Underneath the global variation of total carriage and co-colonization, lie the questions of serotype variation [Imöhl et al., 2010, Moore, 2009]. Why do more than 90 pneumococcus serotypes and strains coexist, and why are particular types present at particular dominant or sub-dominant frequencies in certain locations? This question has intrigued epidemiologists and modelers of pneumococcus dynamics for a long time, especially after the introduction of PCV vaccines. While it is known that each serotype is defined by a unique polysaccharide capsule [Bentley et al., 2006, Geno et al., 2015], hence presenting slightly different within-host growth properties and potentially-varying antigenic properties, it remains still unclear how and to what extent such micro-scale differences translate to higher-level phenotypes to mediate population coexistence [Weinberger et al., 2009], furthermore when compounded with variation in other life-history traits of the microorganism itself, or of the host population [Cobey et al., 2017] - a question not unique to pneumococcus [Koelle et al., 2006, Wikramaratna et al., 2015].

When considering parametrization of pneumococcus phenotypic diversity we must inevitably face the conundrum of how universal these parameters really are, i.e. how much do they vary across settings, and what are the drivers and implications of such variation? Can we, and if yes, how, use the serotype interaction parameters estimated in one context, to understand or predict epidemiology in another? Furthermore, even when assuming consensus on the parameters, the question of how those values impact on the quality of coexistence dynamics of pneumococcus serotypes, their hierarchies, the number and identities of strains composing a given system, the stability and complexity of epidemiological equilibria in a given setting, still remains largely unexplored. The literature is replete with descriptive reports of serotype frequencies, and distribution variation (e.g. see references in [Dekaj and Gjini, 2024] and recent reports [Garcia Quesada et al., 2021, Xu et al., 2024]). However, such empirical picture has not been complemented with a theoretical underpinning based on which such data can be interpreted mechanistically and quantitatively across sites.

Here, we propose a model-based method to tackle this problem. Our approach builds on a general SIS colonization model derived for microbial ecosystems such as endemic pneumococci [Madec and Gjini, 2020], which uses time-scale separation and similarity between strains to simplify coexistence dynamics. This framework arrives down to a very simple replicator equation for strain frequencies over time, expressed in terms of pairwise invasion fitnesses. Although the first link of this N-strain SIS coinfection model with the replicator equation was uncovered in [Madec and Gjini, 2020], it was later extended and derived also in [Le et al., 2023, 2022] allowing for strain variation along 5 traits, and in [Le and Madec, 2025], embedding the epidemiological dynamics in space, hence affording to this link broad generality and wide applicability, beyond the original formulation.

In the following, we show how even this original simple model, with evolutionary-game theory and adaptive dynamics at its core, can be used to integrate context-dependent mechanisms of pneumococcus coexistence across systems, and applied for inter-serotype interaction parameter estimation.

Beyond pneumococcus, this estimation method suggests an avenue for similar applications also in other multi-site diversity data.

## 2 Method

### 2.1 SIS dynamics with coinfection and *N* co-circulating strains

The transmission of *N* strains, follows at its core *Susceptible-Infected-Susceptible* dynamics, with coinfection, described in detail in [Madec and Gjini, 2020], later extended in [Le et al., 2023]. The variables are: *S, I*_*i*_ and *I*_*ij*_, referring respectively to the proportion of susceptible hosts, those singly-colonized by strain *i* and those co-colonized by strains *i* and *j*. How these variables change over time is given by a system of ordinary differential equations. We follow here the original model in [Madec and Gjini, 2020] which is simpler, because it assumes only variation in susceptibilities to co-colonization between strains (*K*_*ij*_):

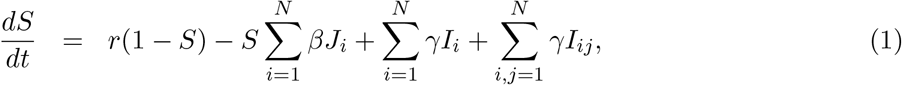

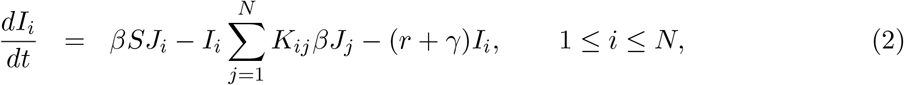

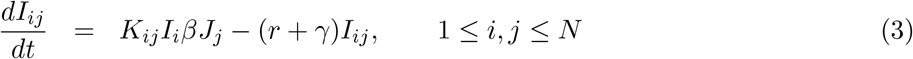

In the above equations, *J*_*i*_ denotes the proportion of all hosts transmitting strain *i*, including singly- and co-colonized hosts, and is given explicitly by 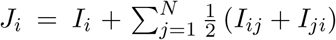. Note that *F*_*i*_ = *βJ*_*i*_ is the force of infection of strain *i*, for all *i* in the system, with *β* representing transmission rate, here assumed equal for all strains. Co-colonized hosts with strain *i* and *j*, transmit either with probability 1/2 (but see [Le et al., 2023] for asymmetric effects). In the system, the parameter *γ* is clearance rate of single and co-colonization episodes, making hosts return to the susceptible class. The recruitment rate of susceptibles is *r*. Susceptible birth rate is assumed equal to natural mortality rate. Upon co-colonization, strains can interact with coefficients *K*_*ij*_: if *K*_*ij*_ *>* 1 this indicates pairwise facilitation (existing strain *i* increases host vulnerability to co-colonization by *j*), while if below 1, *K*_*ij*_ reflect competition in cocolonization between *i* and *j*. The model thus mirrors that a prior colonization by strain *i* alters the susceptibility of the host to co-colonization by strain *j*, without pre-defining the underlying mechanism. Notice that we incorporate in the model co-colonization by the same strain (hence *K*_*jj*_), following theoretical arguments for coinfection model well-posedness and evolutionary neutrality [van Baalen and Sabelis, 1995, Alizon, 2013, Lipsitch et al., 2009].

### 2.2 Strain dynamics at the population level and the replicator equation

By assuming strain similarity, when considering *K*_*ij*_ variation, each co-colonization vulnerability coefficient can be expressed as a benchmark value *k*, plus a small deviation from it:

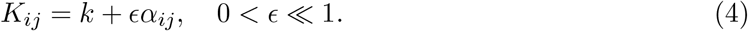

Although there are many choices possible for such representation, as long as *ϵ* remains small, one straightforward choice is for *k* to be the mean of the *K* matrix and *ϵ* the standard deviation. In this case *α*_*ij*_ could be seen as rescaled standardized interaction coefficient between any two strains:

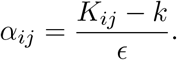

Under this representation, we can apply a slow-fast timescale decomposition to the dynamics, obtaining a suitable model reduction [Madec and Gjini, 2020].

In this model reduction, we find that the fast variables hidden within the system (Eqs. 1-3) are the total prevalence of susceptible hosts *S*(*t*), singly-colonized hosts *I*(*t*), and co-colonized hosts *D*(*t*), which reach their equilibrium (*S*^*^, *I*^*^, *D*^*^) on a timescale *o*(1*/ϵ*):

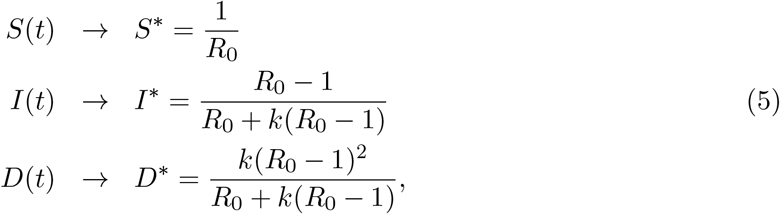

where *R*_0_ = *β/*(*r* + *γ*) denotes the basic reproduction number, equal among all strains [Madec and Gjini, 2020], and *k* refers to the uniform susceptibility to co-colonization, obtained in the limit *ϵ* → 0.

On the other hand, we find that the slow variables are strain frequencies, defined as

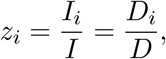

where 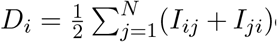 denotes the collective proportion of all co-colonized hosts carrying strain *i*. These strain frequencies follow an explicit replicator equation over the timescale *τ* = *ϵt*:

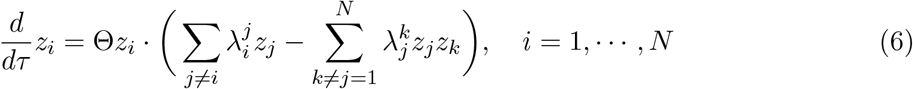

where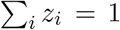, and 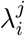 denote pairwise invasion fitnesses (pairwise invasibility coefficients) between any two strains, Θ being the speed of the dynamics.

When strains vary in *only* co-colonization vulnerabilities [Madec and Gjini, 2020], the pairwise invasion fitness between strains *i* and *j* in our SIS co-colonization model, is given by:

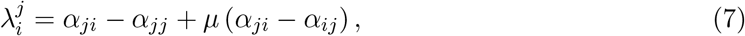

where the quantity

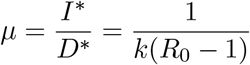

describes the ratio of single- to co-colonization (single- to coinfection) in the epidemiological system.

A remarkable feature of the replicator equation is that it becomes explicit how the interstrain dynamics in the system are coupled and dependent on each other. Each strain grows in frequency depending on how it ‘invades’ other strains in the system (the first summation term in the parentheses in Eq. 6 effectively gives the mean fitness of strain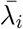), and also depending on how large or small is the mean invasion fitness of the system as a whole (the quadratic term is effectively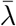). In fact, a strain will only increase in frequency whenever its own invasion fitness exceeds the mean invasion fitness of the system, i.e. when 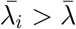 and vice-versa.

Finally, in this slow-fast dynamics model approximation, the prevalences of infected host compartments are obtained from the replicator variables directly as:

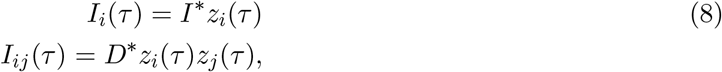

reducing the model computational complexity from quadratic (*N* ^2^) to linear (*N*) in the number of strains, as we do not need to solve the full epidemiological model equations (Eqs. 1-3), but rather just the replicator system (Eq. 6) and then merge the **z** solution with the global aggregated variables (*S, I, D*).

We have shown that due to different biological details, the emergent structure of the invasion fitness matrix 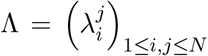 can vary and shape accordingly coexistence dynamics in the multi-strain system [Madec and Gjini, 2020, Gjini and Madec, 2021, 2023]; many regimes being in line with expected outcomes from classical replicator dynamics [Allesina and Levine, 2011, Yoshino et al., 2008, Hofbauer and Sigmund, 1998].

One regime of particular interest is in the limit of *µ* → 0 and *µ* → ∞ for fixed *α*_*ij*_ ~ 𝒩 (0, *σ*^2^): we showed that in the first limit, regimes of (multi-) stable coexistence of a few serotypes occur, whereas in the second limit, a complex, typically unstable, oscillatory dynamics among many serotypes, is the rule [Gjini and Madec, 2021]. This result stems from the way pairwise invasion fitness in this model 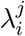 is an explicit function of *µ*, making the ratio of single to co-colonization, a key global epidemiological contextual parameter, able to qualitatively tune selection.

#### Pneumococcus serotype frequency data

The replicator equation is an explicit model for evolution of strain frequencies over time. To link it with data, we considered serotype frequency data reported in epidemiological cross-sectional surveys of *Streptococcus pneumoniae* in the literature. We use 5 reports from different countries, which quantified also global colonization and co-colonization prevalences, analyzed by us in a previous study [Dekaj and Gjini, 2024].

Although the previous study showed that basic reproduction number *R*_0_ and mean cocolonization vulnerability *k* inversely co-vary globally to give rise to a generally conserved ratio *µ* around 6, individual geographic sites do show some empirical variation, which may be relevant, and which was quantified broadly as *low* vs. *high µ* settings.

Our dataset here contains pneumococcus serotype frequency data from both high and low *µ* settings, namely from five geographic settings (see Figure 1): Denmark Harboe et al. [2012], Iran Tabatabaei et al. [2014], Brazil Rodrigues et al. [2017], Nepal Kandasamy et al. [2015], and Mozambique Adebanjo et al. [2018], with ratios of single-to-colonization between 0.93 (Iran) and 16.8 (Mozambique). These settings differ in the number and identities of serotypes reported, as well as their frequencies, however they display a remarkably similar rank-order-abundance distribution (Fig. 1). Furthermore, a number of seven serotypes appear in all 5 settings, these being 14, 19A, 19F, 3, 4, 6A, 6B. The data-files are provided in the Supplementary Materials.

**Figure 1:**
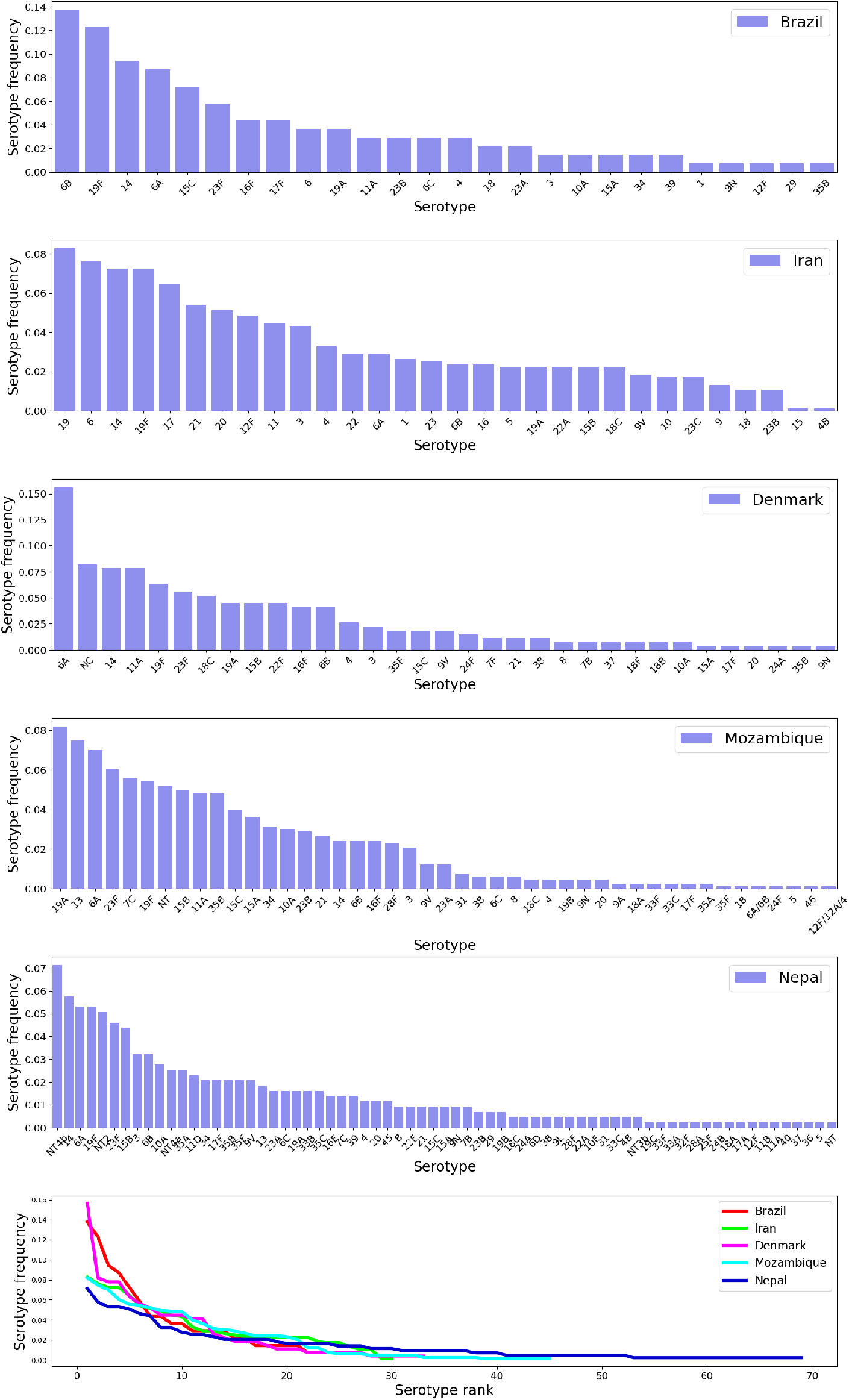
Rank-abundance data for pneumococcus serotype frequencies across 5 geographical sites, presented in the order of increasing number of serotypes. The cross-sectional data are reported in the following epidemiological studies conducted in children populations: Brazil Rodrigues et al. [2017], Iran Tabatabaei et al. [2014], Denmark Harboe et al. [2012], Mozambique Adebanjo et al. [2018] and Nepal Kandasamy et al. [2015]. See also Table 1 for more details.

#### Context-dependence model for invasion fitness parameters between serotypes

The case of bottom-up constitution of the pairwise invasion fitness matrix Λ via random normally-distributed *α* coefficients centered at zero has been explored in depth in [Gjini and Madec, 2021]. This bottom-up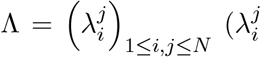’s are defined in (7)) structure can be obtained via fixing the *α*_*ij*_ coefficient probability distribution, and possibly varying the *‘global context’* across settings,

**Table 1:** *S. pneumoniae* in five eco-epidemiological contexts with different global colonization and co-colonization parameters. Here we report mean-field parameters of *Streptococcus pneumoniae* endemic prevalence across 5 studies in children populations (*<* 6 years), estimated by Dekaj and Gjini [2024] using the SIS model with co-colonization [Madec and Gjini, 2020]. The order of presentation in this table follows the increasing number of serotypes reported.

| *Location / Study site* | *Single to cocolo-*<br>*nization ratio* | *Basic reproduc-*<br>*tion number* | *Mean suscep-*<br>*tibility to co-*<br>*colonization* | *Epidemiological*<br>*data* |
| --- | --- | --- | --- | --- |
| (Nr.serotypes/Sample<br>size) | $\hat{\mu}$ (95% CI) | $\hat{R}_0$ (95% CI) | $\hat{k}$ (95% CI) | reference |
| **Brazil** (26/242) | 9.85 (4.09-13.93) | 2.4 (2.28-2.86) | 0.07 (0.06-0.13) | Rodrigues<br>et al. [2017] |
| **Iran** (30/1291) | 0.93 (0.66-1.06) | 1.52 (1.5-1.58) | 2.06 (1.9-2.6) | Tabatabaei<br>et al. [2014] |
| **Denmark** (33/437) | 10.23 (5.44-13.16) | 2.3 (2.22-2.56) | 0.08 (0.06-0.12) | Harboe<br>et al. [2012] |
| **Mozambique**<br>(45/927) | 16.8 (10.05-20.38) | 6.44 (6.14-7.69) | 0.01 (0.009-0.015) | Adebanjo<br>et al. [2018] |
| **Nepal** (69/600) | 3.95 (2.49-4.71) | 1.98 (1.92-2.14) | 0.26 (0.23-0.35) | Kandasamy<br>et al. [2015] |

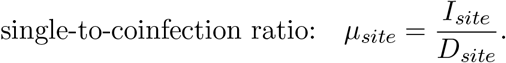

Indeed, this mechanistic formulation can interpolate between regimes of stable coexistence and oscillatory dynamics among serotypes, in the limits of *µ* → 0 and *µ* → ∞, respectively, when *α*_*ij*_ ~ N 𝒩 (0, *σ*^2^).

As this Λ matrix structure encodes context-dependence explicitly, we will consider it as the plausible governing mode of mutual invasibility among multiple pneumococcus serotypes, and without loss of generality, since many different types of replicator dynamics can be captured by this formulation [Gjini and Madec, 2021]. Thus we will assume that, on one hand, the *α* matrix is fixed (universally conserved):

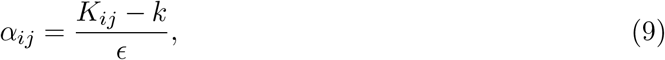

and on the other, the Λ matrix may be different in each system, and can be obtained by substituting the site-specific value of *µ* (Table 1) into Eq.7. Thus, pairwise invasion fitness matrix in each setting is given by:

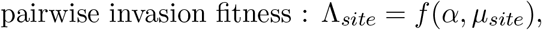

applying the model where strains vary just in vulnerabilities to co-colonization (Eq. 7), while being equivalent in other traits [Gjini and Madec, 2021]. This is a quite general and flexible representation because mathematically it has the capability to generate all types of complex replicator dynamics. Our goal will be to estimate the matrix of *α*_*ij*_ in a way that is consistent with serotype composition and frequency data, and *µ* values across all countries.

#### Equilibrium or non-equilibrium assumption?

We do not have multiple time-points from the same epidemiological setting. Instead we have a snapshot of cross-sectional prevalences from each site. How much can we infer from single crosssectional patterns of serotype frequency variation across settings? The first modeling choice will be how to interpret data in each system, as an equilibrium representation or as a more arbitrary snapshot of the dynamics. To answer this question, we resorted to the most-likely scenario predicted by the model for that empirical *µ* value (Figure 5 in [Gjini and Madec, 2021] for *N* = 10).

According to the values estimated for these 5 geographical settings, we reasoned that Brazil, Denmark and Mozambique having high values of *µ* above 10 on average, are more likely to display non-equilibrium dynamics (*Dyn*), whereas Nepal and Iran, having lower values of *µ* are more likely to display a steady-state scenario (*SS*). This means that these data should be interpreted under these different assumptions, and hence fitted via optimization of slightly different functions. Besides considering whether frequency data reflect a steady state or dynamic snapshot, we must also make a decision in the former case, whether we must impose the strict stability property on the steady-state. We chose to impose the stability criterion whenever a steady-state was assumed. Thus, in our fitting we filtered only those estimates, where the Jacobian of the replicator evaluated at that fixed point had all eigenvalues with negative real part.

### 2.3 Estimating pairwise strain interactions in co-colonization vulnerabilities

We now describe the process estimating parameters, entries of the *α* matrix (*α*_*ij*_) in detail. There are some conditions:

1. We assume strain variations are only in co-colonization interactions: *K*_*ij*_ vulnerability coefficients (sufficient for enabling all regimes of replicator dynamics).
2. We assume all the strains observed in the study were present at equal frequencies at some unknown initial state in the past (in the Dyn assumption cases).
3. We assume the standardized co-colonization vulnerability coefficients of the same strain pair are the same in all the countries 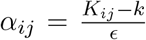 for all (*i, j*) pairs co-occurring across sites.
4. Once the *α*_*ij*_ is estimated for one pair of serotypes, it stays fixed at that value for all other subsequent sites in the fitting procedure.
5. The range of *α*_*ij*_ is fixed to be [−10, 10] across all studies.
6. Not all pairs of strains will co-occur in our dataset, so for the unobserved serotype pairs *i, j* the associated coefficient *α*_*ij*_ cannot be estimated and will remain unknown.

#### Sequential site ordering for the nested *α*_*ij*_ estimation

Our goal is to estimate the matrix of *α*_*ij*_ in a way that is consistent with serotype frequency data **z** and *µ* values across all systems. We adopt 2 ways of ordering the sites of pneumococcus serotype frequency observations: i) one which is just purely based on the increasing number of strains among sites, and ii) another one, which prioritizes stability for the site with plausible stable coexistence between the largest number of strains (see Figures 2-3 for an illustration).

**Figure 2:**
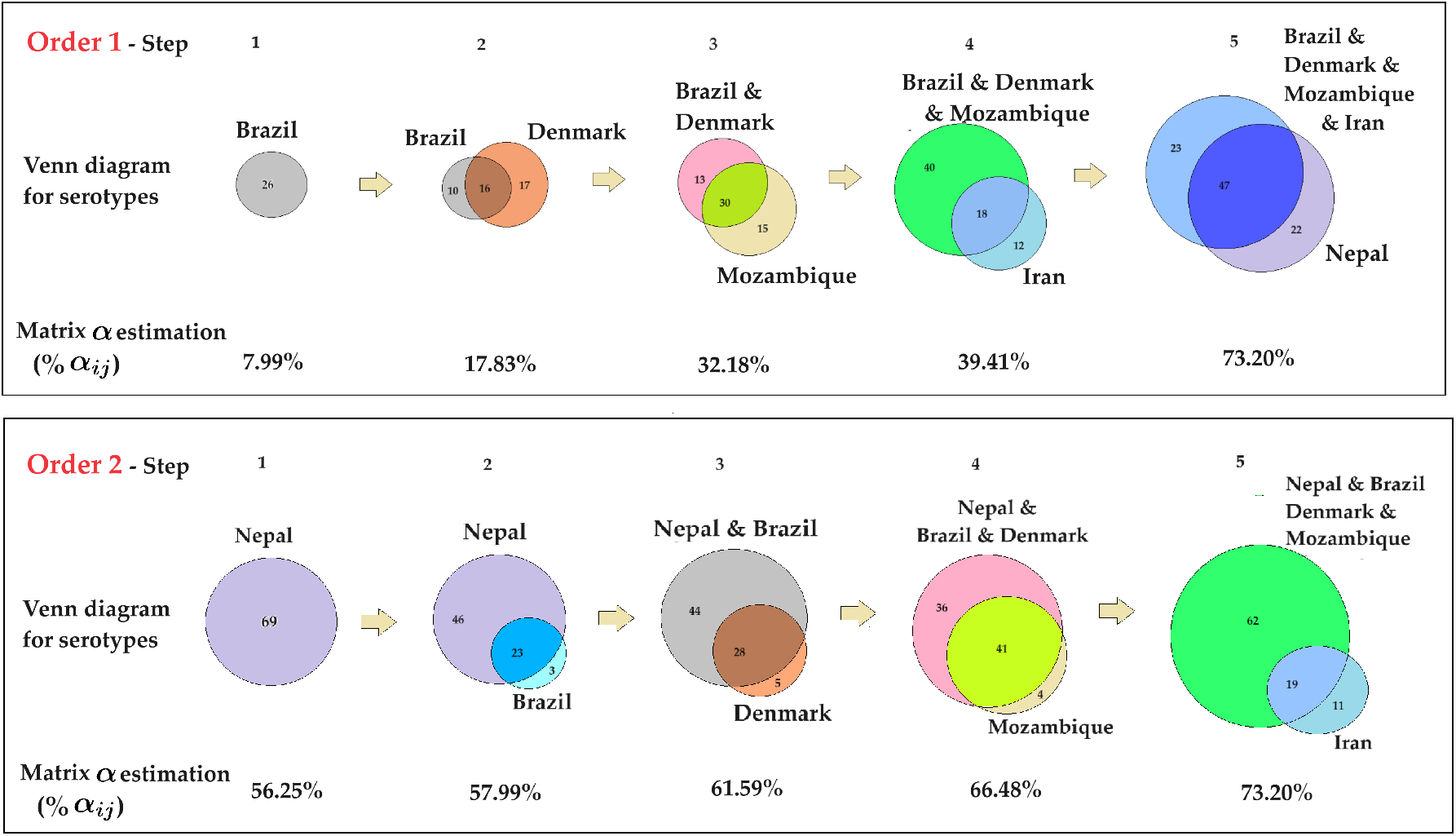
Venn diagram representation of parameter estimation process. We visualize the two approaches to parameter estimation using Venn diagrams. At each step, we highlight the key elements, including the countries with the number of strains already included from previous steps, the country and the number of strains being added in the current step, and the number of common strains between them. Additionally, we calculate the percentage of the 92 × 92 *α*_*ij*_ matrix filled at each step.

**Figure 3:**
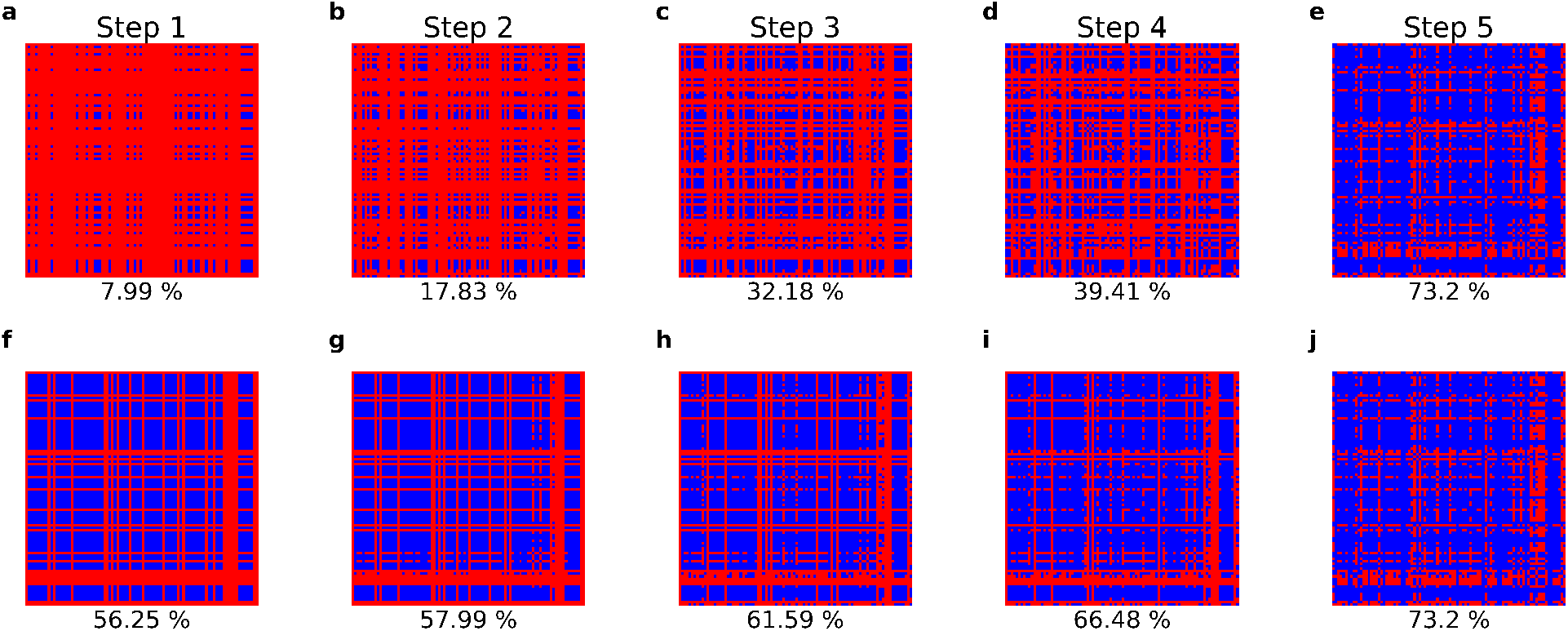
Stepwise filling of the 92 × 92 *α*_*ij*_ matrix in the orders of parameter estimation: Order 1 (a-e), Order 2 (f-j). We further investigate the estimation process by examining the stepby-step filling of the 92 × 92 *α*_*ij*_ matrix. In this visualization, red indicates the *α*_*ij*_ parameters that have not yet been estimated, while blue indicates those that have been estimated up to the current step (cumulative percentages below the matrix). The upper panel illustrates the process for Order 1, while the lower panel depicts the process for Order 2. This matrix-block visualization is consistent with the Venn diagram in Figure 2.

#### Order 1: Site-ordering by “increasing strain number” (lowest → highest number of strains)

The process is carried in a particular optimized order: from Brazil → Denmark → Mozambique→ Iran → Nepal, respectively, based on their number of empirically-observed strains *n*, starting from the site with smallest number of reported serotypes. In this ordering, the first 3 countries are assumed to represent a snapshot of the dynamics, whereas for Iran and Nepal (case 1) a stable steady state is more plausible. Since for Nepal *µ* = 3.95, we also considered the possibility of data reflecting also a snapshot of the dynamics (case 2) but this fit gave higher error. For details please see Supplementary Material S3. In this main part, we use Nepal instead of Nepal-Case 1 for brevity.

#### Order 2: Site-ordering by “stability-of-largest-coexistence” first, increasing strainnumber second

Here the data are fitted in the following order: Nepal→ Brazil→ Denmark→ Mozambique → Iran, with Nepal data being fitted before all others, as a stable coexistence equilibrium of 69 serotypes. We estimate the parameters for Nepal first, followed by the other countries. This approach is necessary because computing the eigenvalues (to verify stable equilibrium) of the Jacobian matrix of Nepal’s fitness matrix (size 69 × 69) is computationally intensive and challenging due to the lower degrees of freedom.

Within remaining countries, the process is carried out in the following order: from Brazil, to Denmark, to Mozambique, Iran, respectively, based on their number of coexisting strains *n*, starting from the site with smallest number of reported serotypes.

In the second case, we re-scale the *α*_*ij*_ variance, immediately after the first country estimation, to match that obtained in Order 1, so that we can compare the resulting interaction parameter distributions between the two methods under the same scale.

For details please see Supplementary Material S4-S5.

#### Error function

In the case of relative abundance data, 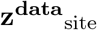 being interpreted as a dynamic snapshot (*τ* = *τ*_0_ = 50), the error for that site (root mean squared error, RMSE) will be given by:

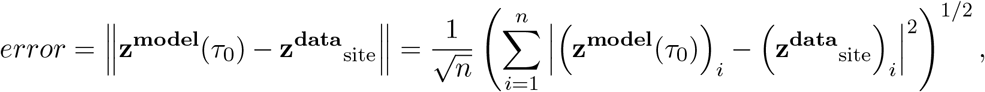

and in the case of assuming 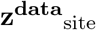 to be an equilibrium (*dz*_*i*_*/dτ* = 0, for all *i*), the error for that site will be given by:

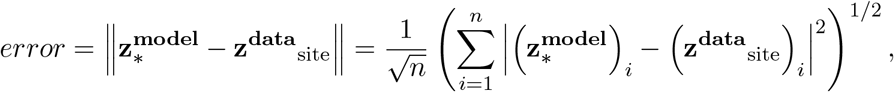

in which, **z**^**model**^ (*τ*) is the predicted frequency of serotypes, i.e. the solution of the replicator dynamics at time *τ*; and 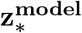 is the stable equilibrium of the dynamics. To denote the first abbreviation case we will use the *Dyn* and for the second case, the abbreviation *SS*. It should also be noted that, in the estimation process, we assume the speed of the dynamics is consistently set to 1. This assumption will be discussed later.

#### The algorithm

In either sequential ordering of our datasets, the algorithm we use in the *α*_*ij*_ estimation process is **Particle Swarm Optimization** (PSO), with the appropriate range in each country (here we use the range [−10, 10] for all sites’ parameters estimation), and starting parameter guess at 0, for the sake of convenience. This choice of range and starting point is valid because adding the same value to all elements of the *α*-matrix does not change the replicator dynamics [Gjini et al., 2016, Le et al., 2023].

PSO is a powerful computational method used mainly for global optimization, that optimizes an objective function by iteratively trying to improve a candidate solution with regard to a given measure of quality [Eberhart and Kennedy, 1995]. Intuitively, each particle’s movement is influenced by its local best-known position and guided toward the global best positions discovered by other particles. This is expected to move the swarm toward the best global solutions. It is different from other optimization algorithms in such a way that only the objective function is needed, and it is not dependent on the gradient or any differential form of the objective. It also has very few hyperparameters, which is an advantage in our case. This optimization algorithm performs well on parameter optimization for ordinary differential equation models [Akman et al., 2018]. However, convergence to the overall optimal solution is not always guaranteed [Raß et al., 2015], due to several reasons, for example, the PSO involving random components in velocity updates; this randomness can lead to different results in different runs and does not guarantee convergence to the global best solution every time. In addition, as the number of dimensions increases, the search space grows exponentially, making it harder for PSO to efficiently navigate and find the global optimum. This is why we decided to fit the data sequentially site by site, and not altogether.

## 3 Results

### Fit and quality of fit

Our algorithm, is able to fit extremely well the observed serotype prevalences in each study site (Figure 4), via both ordering methods, mean error being less than 1% for each ordering of data (Tables S1-S2). While the nested method based on order 1 is less conservative in terms of dynamics and fits the data in the order of increasing number of serotypes, the second method based on order 2, fits first the largest number of strains adopting the stability constraint. This means the second method constrains a larger proportion of the alpha matrix from the beginning, an effect that trickles down to all subsequent site fittings. We can see that the estimates obtained by order 2 yield generally lower errors than those obtained with order 1.

**Figure 4:**
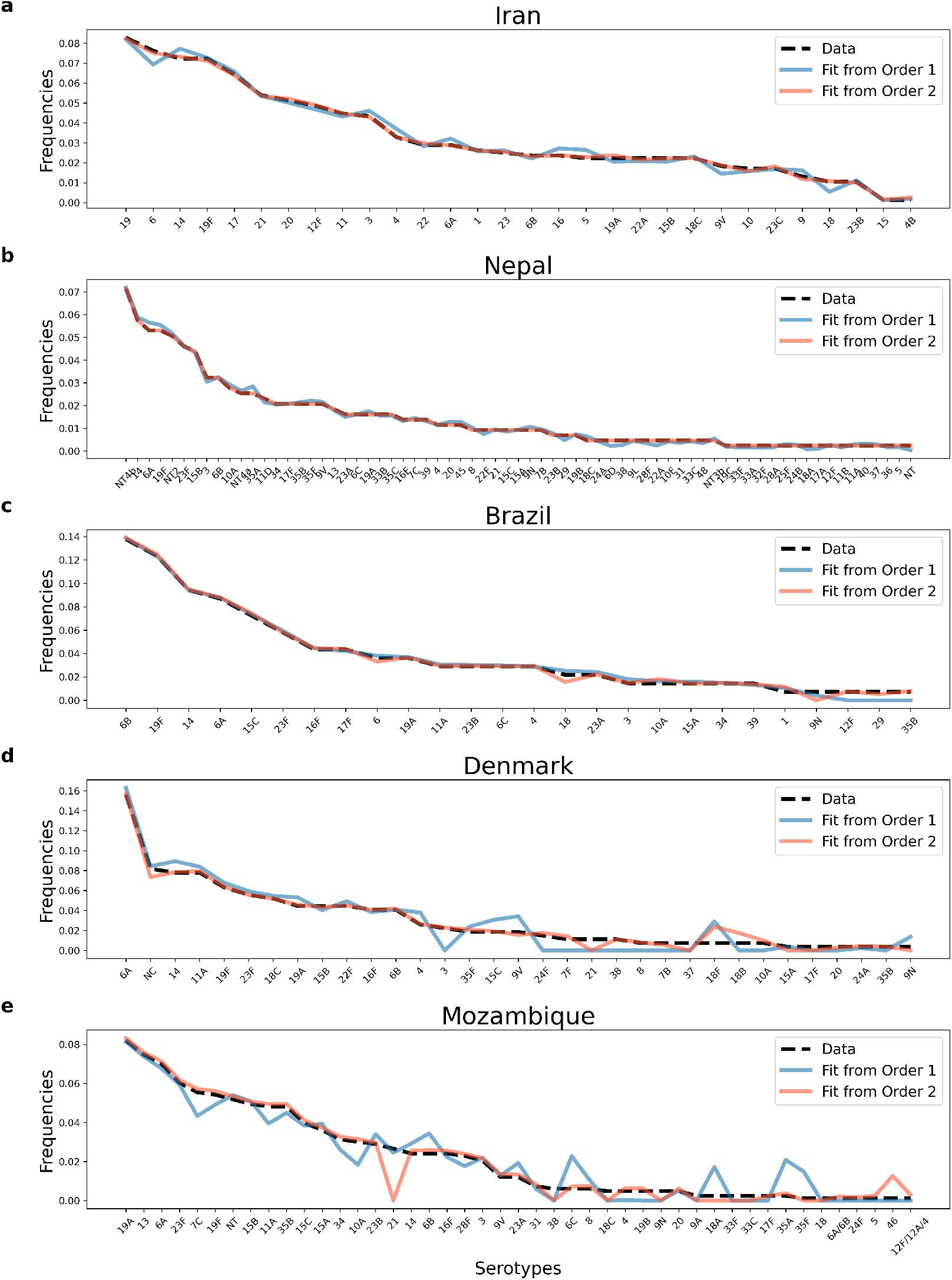
Relative abundance data and model fits for pneumococcus serotype frequencies in 5 epidemiological contexts (ordered by ratio of single to co-colonization, from lowest to highest). We plot the data extracted from Denmark Harboe et al. [2012], Iran Tabatabaei et al. [2014], Brazil Rodrigues et al. [2017], Nepal Kandasamy et al. [2015], and Mozambique Adebanjo et al. [2018], as well as the model-predicted relative abundances (estimated from nested data fitting in Order 1 and Order 2), ranked from the most prevalent to the least prevalent serotype. For details on quality of fits see Tables S1-S2.

When computing the proportion of predicted serotype frequency data within a margin of 10% error for all countries from both orders (Figure S1), we find that in Order 1, the percentage of serotypes predicted within an accuracy of 10% are: Iran 73.33%, Nepal 50.72%, Brazil 65.38%, Denmark 30.3%, and Mozambique 33.33%. Meanwhile, in Order 2, these percentages are higher: Iran 93.33%, Nepal 100.0%, Brazil 80.77%, Denmark 51.52%, and Mozambique 46.67%. The lower prediction accuracy for Denmark and Mozambique is due to two reasons: i) in order 1, the main reason is their dynamics is fitted as a snapshot solution of the replicator, and ii) in order 2, the degrees of freedom in their serotype sets are lower. This means most serotypeinteractions are already estimated in the preceding sequential procedure. For example, in order 2, Mozambique had only 4 over 45 serotypes free to inform the *α*_*ij*_ estimation.

### Statistics of the estimated random α matrix

As shown in Figure 5, the distributions of the standardized co-colonization coefficients between serotypes obtained by either nested method have means near 0 and standard deviation near 1. The distribution of *α*_*ij*_ in Figure 5a is nearly symmetric, meanwhile in Figure 5b, it is left-skewed, as we can see the outliers near −10. There are 73 outliers with values below −7; 69/73 of them belong to the diagonal 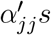 from Nepal *α*-matrix. Compared to the kurtosis of normal distribution, which equals 3, both of these distributions have heavier tails, i.e. are leptokurtic. Especially, the distribution in Figure 5b has extremely thin tail, due to the much more dense concentration of *α*_*ij*_’s around 0. The estimated set of *α*_*ij*_ around the negative value −10, arises as a result of imposing the stability criterion on a large number of strains (69 serotypes in Nepal) from the beginning. The fact of these negative coefficients coinciding with diagonal entries of the *α* matrix, i.e. within-serotype interactions, is consistent with stabilizing mechanisms of complex multi-species coexistence [Allesina and Tang, 2012].

### The mutual invasion fitness network that emerges

The pairwise invasion fitness matrix, which is the higher-order fitness modulated by contextual parameters such as the ratio of single and co-colonization, appears to be highly anti-symmetric for pneumococcus serotypes (dominated by +− edges) under the estimation via order 1, and more skewed towards the pairwise coexistence region (dominated by ++ edges) under the estimation via order 2 (Figure 6 and Figure 7). This means that when imposing few stability constraints for the collective coexistence of subsets of serotypes, the system is more free to vary, and parameter values compatible with observations, reflect more often ‘prey-predator’ mutual invasion outcomes leading to an oscillatory complex coexistence between serotypes, and more rarely a stable one [Gjini and Madec, 2021].

**Figure 5:**
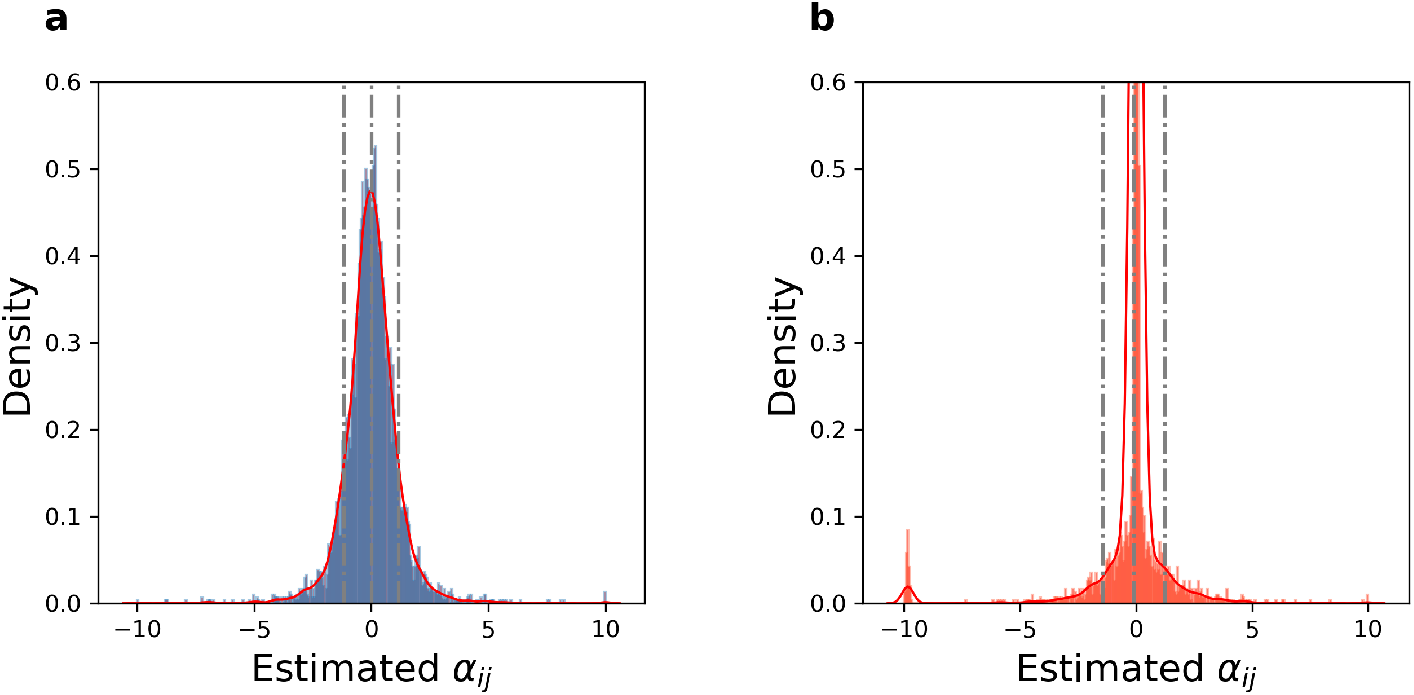
Final global *α* matrix distribution of rescaled interaction coefficients (*α*_*ij*_) for observed serotype pairs across 5 epidemiological contexts of Streptococcus pneumoniae colonization, after the nested fitting of cross-sectional data in Order 1 and Order 2. The figures illustrate distributions of the all *α*_*ij*_’s (73% of the entire matrix A) estimated in the two estimation approaches. We note that both distributions appear bell-shaped, although the second case is much more highly peaked at 0. The statistics for the two distributions are as follows. **(a)**- Order 1: Mean: 0.0038, Standard Deviation: 1.1610, Skewness: 0.1862, and Kurtosis: 13.2259. **(b)**- Order 2: Mean: −0.0973, Standard Deviation: 1.3389, Skewness: −4.0790, and Kurtosis: 37.8318. For site-specific distributions see Figure S2.

**Figure 6:**
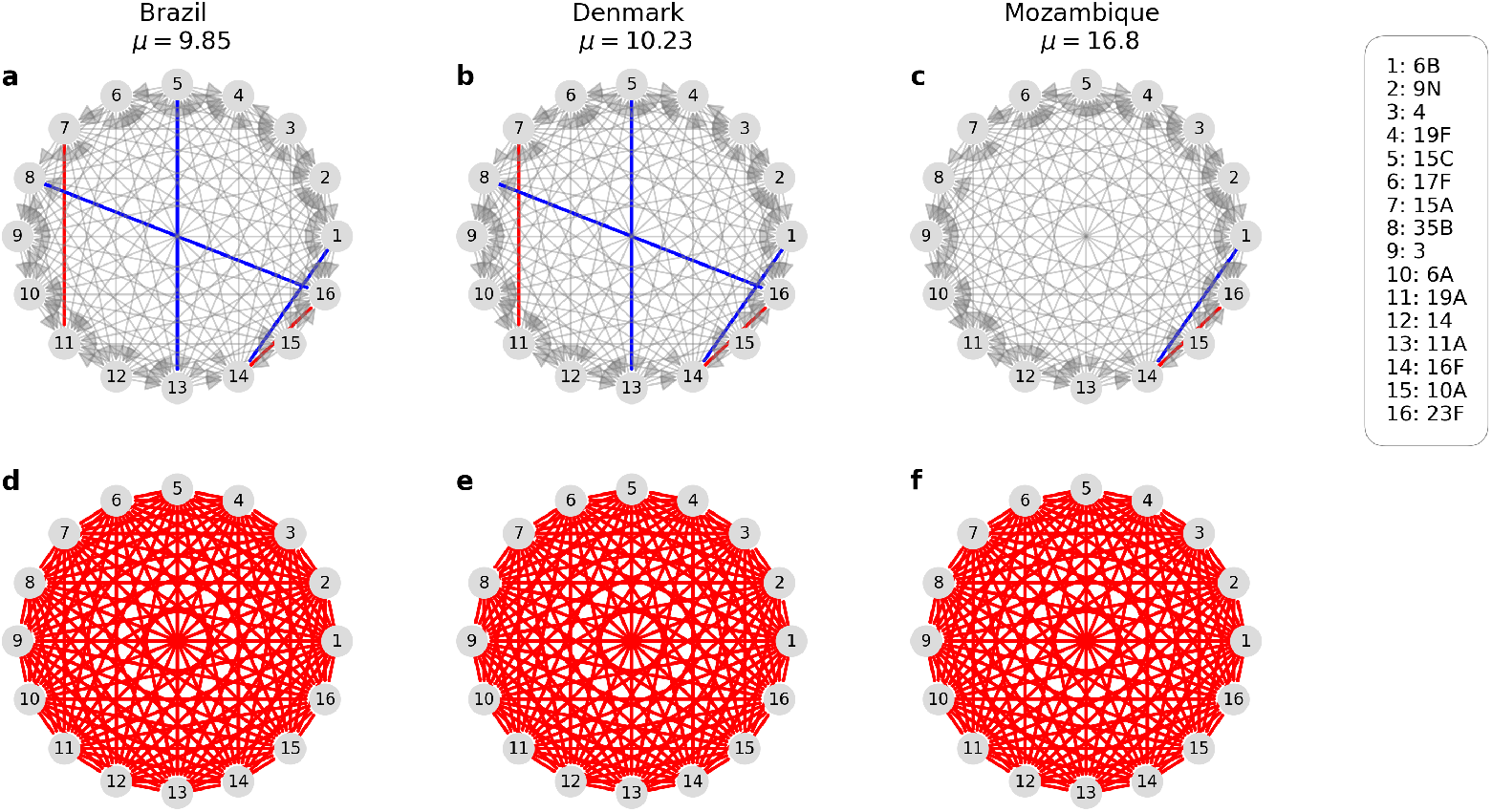
Emergent invasion fitness network among 16 pneumococcus serotypes in different epidemiological settings. We show here the obtained mutual invasion fitness network for the cooccurring serotypes in countries where a dynamic snapshot of ecological trajectories is assumed (sub-part includes only common serotypes). The upper panel visualizes the network from Order 1, dominated by +− edges (gray arrows) with rare coexistence (red) or bistability (blue), meanwhile, the lower panel represents the network obtained from Order 2 data fitting, constituted entirely by pairwise coexistence edges ++ (red lines) for these 16 serotypes.

**Figure 7:**
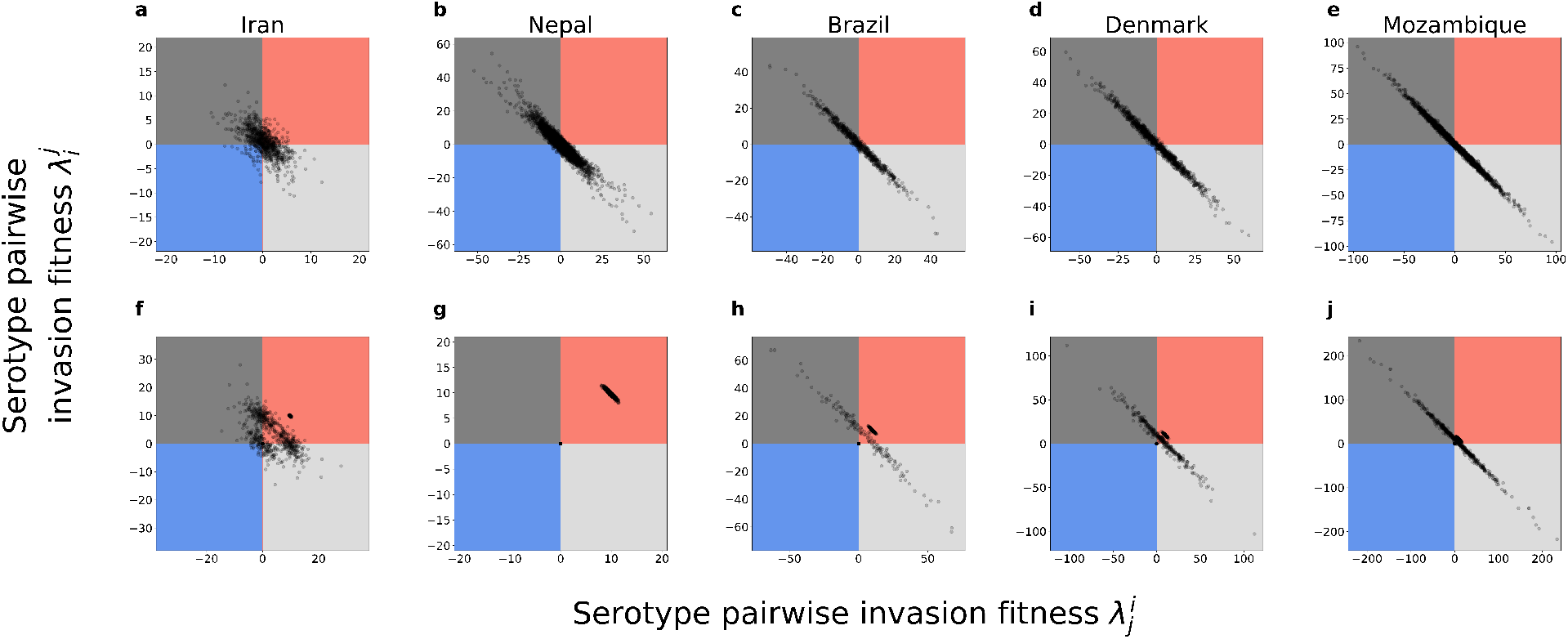
Quantifying coexistence patterns of different pneumococcus serotype pairs across 5 countries resulted from two approaches. We plot the resulting pairwise invasion fitnesses between all serotypes present in each study site. **Top row:** obtained by nested method of order 1. **Bottom row:** obtained by nested method of order 2. For more details, we compute the percentages of pairwise outcomes (edges of the network), including exclusion of either strain, bistable outcome, and coexistence, for each country and each estimation order. We report the sequence triple (exclusion of either strain, bistability, coexistence) as the percentage among all pairwise outcomes. We have that, in Order 1: Iran (69.8%, 7.6%, 19.3%), Nepal (88.2%, 4.5%, 5.8%), Brazil (91.7%, 2.4%, 2.1%), Denmark (94.0%, 1.7%, 1.3%), Mozambique (95.0%, 1.3%, 1.5%). Meanwhile, in Order 2: Iran (41.8%, 2.9%, 52.0%), Nepal (0.0%, 0.0%, 98.6%), Brazil (17.5%, 0.0%, 78.7%), Denmark (21.1%, 0.0%, 75.8%), Mozambique (22.3%, 0.0%, 75.5%). This confirms that the second nested approach yields relatively many more coexistence pairwise outcomes.

In contrast, when we start the estimation by imposing stable coexistence first among a large collective of serotypes (Order 2) then, the parameter estimation is more prone to converge towards values compatible with a majority of serotype pairs in the mutual coexistence region (++ edges), leading to stable coexistence everywhere, independently of contextual parameters or local serotype composition.

Next, we use the estimated invasion fitness matrix to compare invasion resistance Q in each pneumococcus multi-strain system (Figure 8). The mean invasion resistance values for all countries, resulting from Order 1-estimation, are significantly lower (all below 1) compared to the *Q* values obtained from Order 2 estimation (all above 7, except for Iran, where the *Q* value just exceeds 5). It is striking that although the numbers and identities of serotypes vary vastly across settings (Figure 1), the final collective invasion resistance of the pneumococcus multi-strain system is quite similar everywhere.

**Figure 8:**
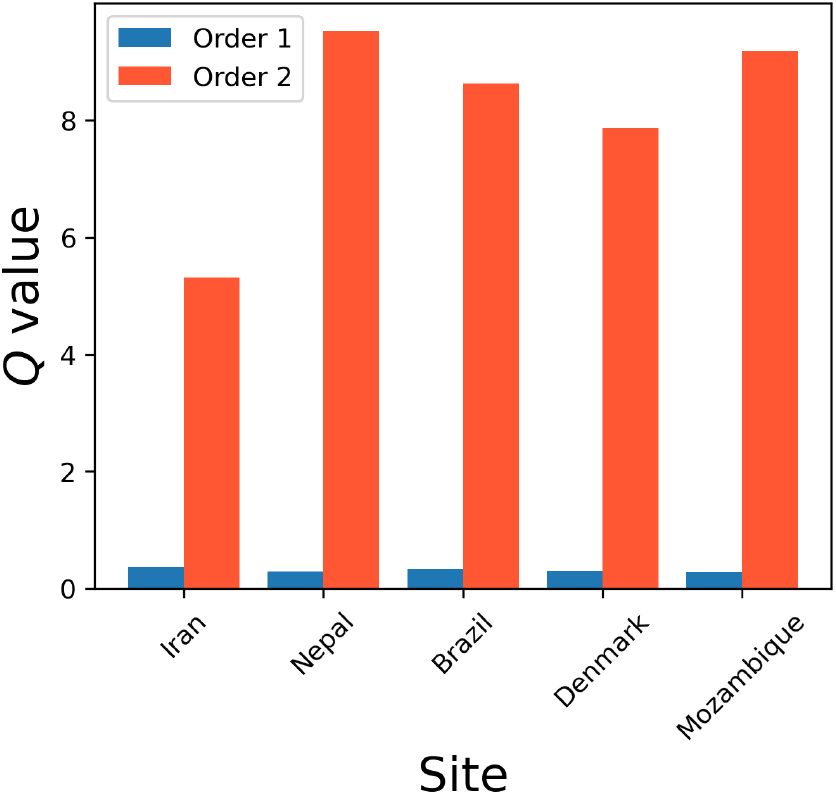
Resulting mean invasion fitness 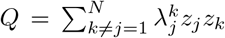 for the multi-serotype pneumococcus ecosystem in each epidemiological site. In each estimation order, we use the finallyestimated *λ*^*′*^s to visualize the 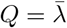 values for Iran, Nepal, Brazil, Denmark, and Mozambique, at the *z* values reported in each setting. Under stable coexistence (predicted from Order 2 multi-site data fitting), it would be harder for new strains to invade, as 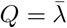 is higher.

### Symmetry of co-colonization interactions (*α*) and invasion fitness (Λ) matrix

The symmetry of the estimated matrices is also important. The symmetry in rescaled interaction coefficients *α*_*ij*_ and pairwise invasion fitnesses 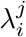 reveals different properties. To explore the sensitivity of the dynamics to *µ* (the ratio *I/D* in each site), we consider the symmetry of the *α* matrix. To understand the local quality of the dynamics, we consider the symmetry of the Λ matrix [Gjini and Madec, 2021]. The level of symmetry of a matrix 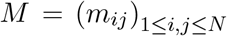 is calculated as described in Tokita [2023]:

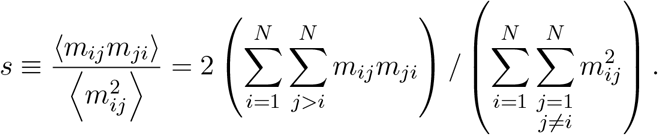

The closer the *s* value is to 1, the more symmetric the matrix is, and conversely, the closer *s* is to −1, the more skew-symmetric the matrix is.

We find the matrix *α* has a symmetry index very close to 0, which indicates a random distribution. This confirms that the dynamics are sensitive to *µ*.

When we consider the 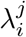 matrix, for each country (Iran, Nepal, Brazil, Denmark, Mozambique), we find that in Order 1, the resulting invasion fitness matrix has a symmetry index very close to −1 in most sites, whereas in order 2 has a symmetry index close to 1. This indicates that under order 1 parameter estimation, the predicted serotype dynamics is likely to be complex and oscillatory, while under order 2 parameter estimation, the dynamics are more likely to be stable coexistence. For details see Table 2.

**Table 2:** Summary of symmetry levels in (*α*_*ij*_) and Λ matrices (≈ 70% of 92 × 92 serotype pairs, cumulated across five countries), after estimation via 2 site orderings. These computations confirm the long-term behaviors depicted in Figure 9. In Order 1, all five countries exhibit a random distribution of *α*_*ij*_ values. A similar pattern is observed in Order 2; however, Nepal’s (*α*_*ij*_) matrix is slightly more anti-symmetric, while Brazil’s matrix is slightly more symmetric. Regarding the symmetry levels of the Λ fitness matrix, in the Order 1 panel, except for Iran and Nepal, where the *µ* values are small enough and data observations are regarded as stable equilibrium, the other three countries—Brazil, Denmark, and Mozambique—display nearly anti-symmetric fitness matrices, which are associated with oscillatory long-term serotype behavior. In contrast, in Order 2, all five countries have random or nearly symmetric Λ fitness matrices, consistent with the convergence to stable states in their dynamics.

| Estimation | Matrix | Iran | Nepal | Brazil | Denmark | Mozambique |
| --- | --- | --- | --- | --- | --- | --- |
| **Order 1** | $(\alpha_{ij})$ | 0.017 | 0.003 | 0.038 | -0.011 | -0.047 |
| | $(\lambda_i^j)$ | -0.61 | -0.96 | -0.99 | -0.99 | -0.99 |
| **Order 2** | $(\alpha_{ij})$ | -0.035 | -0.188 | 0.164 | -0.006 | 0.003 |
| | $(\lambda_i^j)$ | 0.34 | 0.99 | -0.12 | -0.34 | -0.79 |

### Site-specific predictions

Finally, we verified that in those countries where we expected oscillatory dynamics, via *µ* gradient arguments [Gjini and Madec, 2021], we obtained oscillatory dynamics in simulations starting from equal relative abundances between serotypes at some point in the past (Figure 9). This was true for the Dyn countries in Method 1 (Brazil, Denmark, Mozambique). We also verified that in Iran and Nepal (method 1), where we imposed a stable equilibrium within the fitting procedure (SS), the estimated *α*-matrix, and then Λ-fitness matrix, generated a dynamics converging to that stable equilibrium when starting close enough (naturally, all eigenvalues had negative real part).

**Figure 9:**
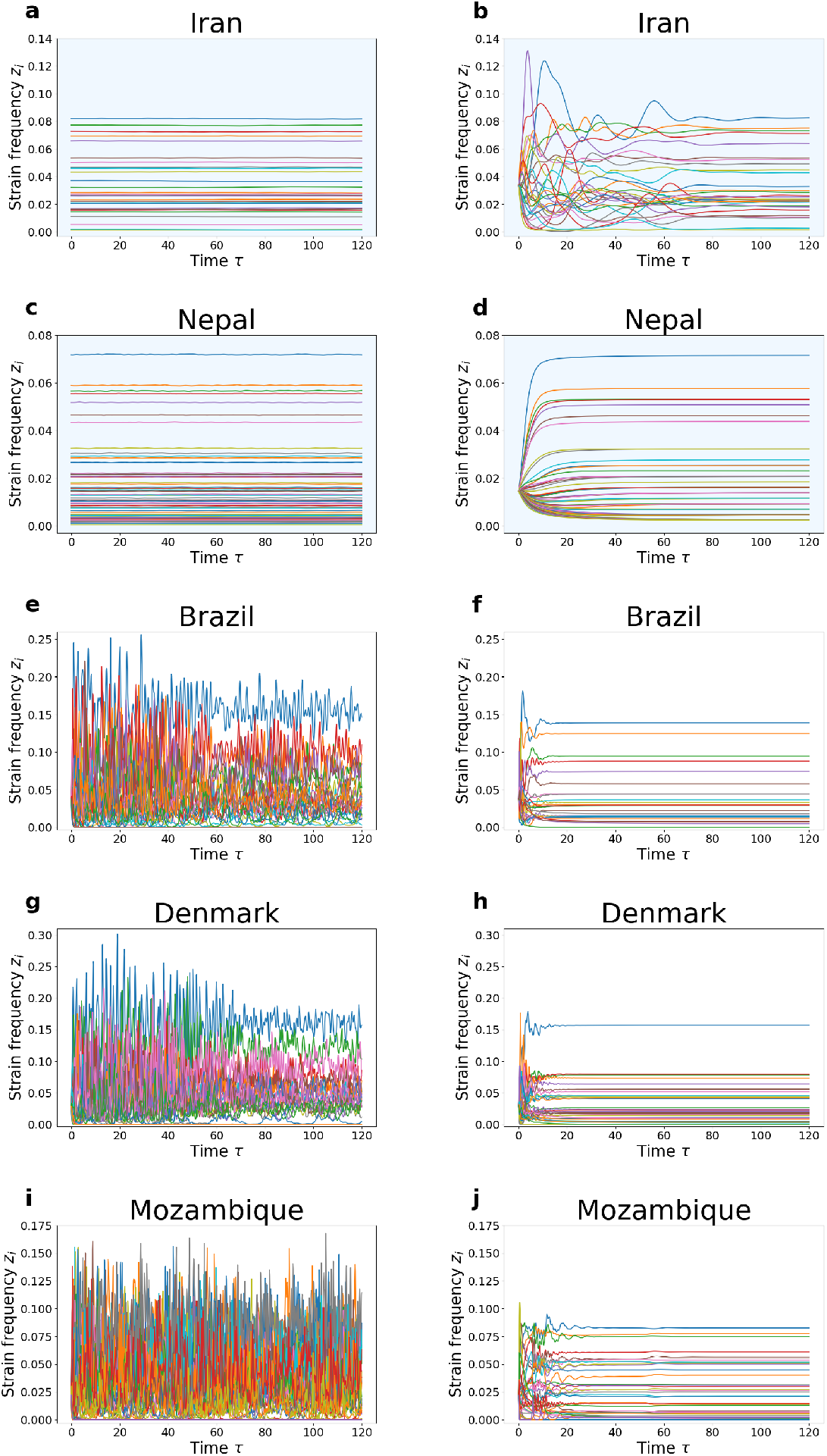
Model predictions for temporal dynamics of multiple strain frequencies at the population level. **Left:** Order 1. **Right:** Order 2 predictions. We llustrate the model-predicted dynamics of serotypes in each geographic site, using the estimated co-colonization and invasion fitness parameters obtained from Order 1 fitting (left) and Order 2 fitting (right). The light-blue shaded background denotes cases where the data are fitted to a strict stable equilibrium (a & c, b & d), while the white background indicates cases where data observations are considered as snapshots at *τ* = 50 of the dynamics. For scenarios where data are fitted as snapshots in both orders, initial conditions assume equal frequencies for all observed serotypes in each setting (e, f, g, h, i, j). In Order 1, where data are fitted to a strict stable equilibrium (a & c) with a narrow basin of attraction, the initial point is set within a small vicinity of the stable state. Conversely, in Order 2 (b & d), the basin of attraction is sufficiently large, allowing the initial conditions to be set at equal frequencies for all observed serotypes while still converging to equilibrium. In this visualization, we assume that the speed of dynamics of all figures are all equal to 1 i.e. Θ = 1.

According to the dynamic simulations in the second column of Figure 9, we observe that in Order 2, the dynamics of the five countries converge to their respective stable equilibria. For Nepal and Iran, where we explicitly impose the condition of a stable equilibrium *a-priori* during the data fitting process, stable coexistence of *all* reported strains is evident (Fig.9 b, d), even when starting from equal relative abundances. Additionally, in the other countries, where data observations are treated as snapshots of the dynamics at *τ* = 50, convergence to a stable coexistence steady state is also confirmed computationally, but with not all observed strains (Fig. 9 f, h, j). Enforcing stability in Nepal results in pushing the diagonal entries *α*_*ii*_ towards negative values, which, in turn, shifts the spectrum to the left half-plane, even after introducing new strains in the subsequent countries. Specifically, the maximum real parts of the eigenvalues of the Jacobian matrices, corresponding to the predicted equilibria of coexisting strains, are all negative, albeit small, as follows: Brazil (25 strains coexisting at the resulting predicted SS, hence 1 observed strain missing from the prediction) −0.06; Denmark (28 strains coexisting at the predicted SS, hence 5 observed strains theoretically extinct) −0.05; and Mozambique (35 strains coexisting at the predicted SS, hence 10 observed strains, theoretically extinct in the simulation), −0.008.

While the existence of alternative stable steady states is interesting to explore, it is indeed computationally very intensive, especially for sites with a large number of strains, and falls outside the immediate scope of this paper. We did this check only for the Iran dataset. When checking the existence of alternative stable steady states for Iran (under parameters estimated via Order 2), we did not retrieve any other steady state under the corresponding estimated Λ-fitness matrix, suggesting the fitted steady state (Fig. 9 b) is indeed a unique monostable equilibrium.

### Reconstructing co-colonization coefficients *K*_*ij*_

How can we obtain the epidemiological co-colonization coefficients *K*_*ij*_ from the estimated *α*’s? We need a value for the small parameter *ϵ* in each site. One way we adopted to get a rough estimate of this parameter was to utilize the uncertainty intervals for *k* (Table 1) and use them to restrict the range of *ϵ* compatible with these bounds. To guarantee universal non-negativitity of each realized 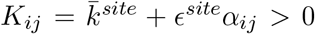 in each site, we must have 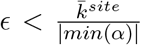, dependent on the ratio between mean *k* and the absolute minimum of *α*_*ij*_ as specified at the outset of our method. In our case, the range of *α*’s was [−10, 10], so a sufficiently low epsilon is approximately *ϵ* = *k/*10, although higher, i.e. less conservative, values are also feasible, depending on the actual estimated *min*(*α*) in each site. Under this scaling we obtained different small values of *ϵ*^*site*^ for the sites modeled here, ranging from 0.007 in Brazil, to 0.2 in Iran, to 0.08 and 0.001, in Denmark and Mozambique to 0.026 in Nepal, indicating slightly different timescales in the slow dynamics in each site.

In conclusion, although recognized theoretically and quantified for many years, co-colonization as an organizing force in pneumococcus has only recently received more modeling and analytic attention in its full high-dimensional structure [Madec and Gjini, 2020, Le et al., 2023]. It is thanks to this mathematical approach, based on similarity arguments, that we can clearly separate the epidemiological dynamics of pneumococci into an explicit fast and slow component, where the *fast* corresponds to globally conserved population-level quantities, and the *slow* corresponds to non-neutral strain dynamics where selective forces (shaped by context) manifest. We believe with the methodological inference framework outlined and illustrated here, it is now time to harness this model’s full empirical potential and predictive power on more diversity data across scales.

## 4 Discussion

Understanding the mode and strength of strain interactions in infectious diseases with multiple co-circulating strains is important, both for interpreting past dynamic trends and for predicting potential ecological and evolutionary changes in the face of vaccination, antibiotics or other pressures [Wikramaratna et al., 2015, Gjini et al., 2016, Alizon et al., 2013, Auranen et al., 2010].

In general, there are 4 types of top-down data-driven approaches for estimating interactions: 1) using longitudinal data in the same host population over time (mostly addressed with Markov models) 2) using cross-sectional data in one host population over time (more easily addressed with dynamic epidemiological models over time), comparable to dynamic gLV models applied to microbiota time-series, 3) fitting one cross-sectional snapshot in one host population (inevitably needs some steady-state assumption for parameter estimation) 4) integrating cross-sectional snapshots in multiple host populations (the most challenging case, addressed here), making use of the largest amount of data and heterogeneities. While the first three above approaches are more represented in the literature, the fourth approach is more rarely found because of all the intricate assumptions for parameter correlations across host/environmental settings that need to be clarified.

### On estimation assumptions and heuristics

Here we believe to have made a significant step in a challenging methodological direction. Our main development is to propose a framework by which multiple strain frequency cross-sectional data from different sites can be integrated under the same model, using context-dependence of fitness parameters and the replicator system.

Although the formal derivation of the replicator system for an SIS co-colonization model was based on the quasi-neutrality (*ϵ* → 0) assumption, in practice we find that the replicator model captures well the co-colonization multi-strain dynamics even for values of *ϵ* = O(10^−1^) (Figure S4 in [Madec and Gjini, 2020]). This means that the validity of our replicator approach should extend beyond the strict similarity condition.

We have highlighted that a key assumption is the order of nesting of parameters in different sites, and this can produce very different parameter estimates, while preserving a very good quality of fit (Figure 4). Also, we find that the way parameters are constrained from the start has a strong bearing on the subsequent quantification of interactions between new serotype pairs. In particular, if we impose the stable coexistence criterion among a large proportion of serotypes from the global pool, this implies strong stability constraints also on the other portions of data fitted in subsequent steps. In our case, this order, stability first, led to better quality of fit of the data, with a global data mean RMSE of 0.0025 (Table S2), while the order 1, in increasing number of serotypes, led to a slightly worse fit, with a global data mean RMSE of 0.0046 (Table S1).

In our fitting procedure, fitting a stable equilibrium with fewer degrees of freedom and more strains is time-consuming (eigenvalues must be computed with negative real parts, and equilibrium must align with observation data). In contrast, fitting dynamic snapshots is easier than a stable equilibrium. There are more degrees of freedom when fitting snapshots. We used a model-informed heuristic for deciding which sites to fit as stable steady states and which sites to fit as dynamic snapshots [Gjini and Madec, 2021]. The ratio of single to co-colonization, *µ*, was used to decide whether we fit a “snapshot” (*µ* relatively large) or a “steady-state” (*µ* relatively small), but here we also imposed on the estimation procedure a strong enforcement of stability. In nesting the data according to Order 1, we prioritized accuracy over stability for Iran and Nepal, where achieving stable equilibrium required considerable effort, especially Nepal with a large number of strains, although eventually we were able to fit it quite well at the end. In the nesting of multi-site data according to Order 2, with uncertainty around Nepal’s data being a snapshot or stable equilibrium, we decided to emphasize stability. Given Nepal’s high number of serotypes, we fitted it first to ensure a stable coexistence equilibrium of all its serotypes (with very little effort). Brazil followed Nepal because fitting any other country before Brazil would have reduced our degrees of freedom to zero, making it impossible to fit Brazil later. Iran, having unique strains, was placed at the end to avoid increasing estimation errors for other oscillating-dynamics countries (Order 2).

A key assumption in our approach was the *µ*-dependence in the pairwise invasion fitnesses, but keeping the essence of each strain pair in terms of fixed *α*_*ij*_. When we repeated the estimation procedure, dropping the context-dependence, and assuming instead the same approximate *µ* ≈ 6 everywhere [Dekaj and Gjini, 2024], and a dynamic snapshot in each site, the fit of the serotype data was slightly worse but the estimated global *α* matrix turned out to be very similar to the one obtained under order 1, with the same pattern resulting in pairwise invasion edges of predator-prey type (Table S3).

Our method is based on the slow-fast dynamic approximation leading to the replicator equation [Madec and Gjini, 2020]. We have chosen to fix the scale of *α*_*ij*_’s in different sites and the time-point of simulating hypothetical dynamics *τ* = 50, starting for equal initial relative abundances. We have also taken Θ, the speed of dynamics in each country, equal to 1. The only variation we have taken into account is *µ* and of course *k* variation (Table 1). This gives freedom to then assign real time units to *τ* according to an appropriate scaling in each country if data from another time-point becomes available. We could indeed use the estimated matrices to make dynamic predictions for later or earlier time points, linking to the year of the data used in our estimation, for example for Denmark, connecting data from [Harboe et al., 2012] to another dataset [Auranen et al., 2010].

### Limitations

Inevitably, our method has limitations. Fitting each site in a sequential manner means that the *α*_*ij*_ is informed by only one epidemiological dataset for each pair of serotypes. Besides the sequential estimation process, we already considered the simultaneous estimation. This approach involves estimating the large *α*-matrix of size 92 × 92, with the objective function being the sum of estimation errors across the five sites, subject to the condition of stable equilibria for Iran and Nepal dynamics (i.e., the Jacobian matrix at equilibrium has all eigenvalues with negative real parts). However, this approach is unfeasible with standard computational resources due to the complexity of the optimization function. Specifically, each iteration requires computing at least three dynamics, two steady states, and the eigenvalues of two matrices of sizes 45 × 45 and 69×69. Another critical challenge arises from the PSO algorithm itself. When the dimensionality increases, the search space grows exponentially, making it difficult for PSO to efficiently locate the global optimum. For these reasons, we opted here for a sequential site-by-site fitting strategy rather than a simultaneous estimation approach, but this remains an avenue for future numerical developments.

We have not taken into account the uncertainty in *µ* or the frequencies of serotypes related to host sample size (Table 1). Instead we have assumed a fixed known ratio *I/D* in each site. We limited ourselves to considering only extreme assumptions about coexistence, either a totally arbitrary dynamics snapshot or stable coexistence fixed point. In general, the replicator system has much more varied dynamics [Gjini and Madec, 2021]. Roughly speaking, when *µ* ≪ 1, there are often several linearly stable stationary states (the system is multi-stable). In this case the dynamics depends strongly on the initial state which is not known here. Note, however, that our estimates do not seem to produce this multistability situation.

By contrast, when *µ* ≫ 1, the attractors are more likely periodic trajectories or even strange attractors. Our methodology does not take into account these constraints. We could relax the assumption of stability in later investigations, including for example permanence of the observed species [Aubin and Sigmund, 1988] as a criterion in the estimation procedure, imposing that the mean frequency over time corresponds to the observed data. Our estimation of the 92 × 92 interactions matrix is not complete. We could only estimate 73% of the entries of this matrix for the pairs of serotypes that were co-observed in the same site at least once. This poses limits to our prediction with the estimates we have got. In other settings where the serotype composition is entirely contained in the estimated portion, we can apply the model to make predictions of the dynamics. In other settings, where new serotype pairs might be observed, we can adopt two approaches: i) we will either have to estimate the new pairwise interactions, keeping fixed those interactions that were already fitted here, or ii) try to make predictions by assigning values to new *α*_*ij*_, e.g. resampling them from the current distribution, or setting them to zero. New sites will provide new data to both test the validity of our obtained estimates on one hand, and to expand the estimation procedure toward the unknown portion on the other.

We also did not consider sub-serotype diversity, assuming to a first-order approximation, homogeneity within serotype in pneumococcus. This is a strong assumption, but other layers of diversity could be explored in the future. The model structure could also be altered or expanded, as we have already mentioned previously [Madec and Gjini, 2020, Le et al., 2022, 2023]. Increasing model complexity however, would possibly augment nonlinearity and computational requirements for parameter estimation. It remains yet an interesting avenue to explore.

Given the limited scope of our study, we also did not examine in detail the identities of serotypes displaying certain interactions and certain invasion fitnesses, but the results of our matrix estimations are all public (see S5), and could be investigated in follow-up studies, to elucidate serotype-specific patterns and signatures of dominance on the collective dynamics.

### Towards bottom-up trait-based interactions and invasion fitness

A critical issue in modelling multi-strain systems is how to best represent the relationships between the biological entities involved. A common approach is to consider strain space as a continuous volume, surface or even a single dimension where the Euclidean distance between points serves as a metric for antigenic and/or genetic similarity [Gog and Swinton, 2002, Lin et al., 1999, Bedford et al., 2012].

We could link our estimated *α*_*ij*_ distribution for pneumococcus serotypes to some genetic or antigenic distance between them, like it is done for influenza models or other more abstract multi-strain models, a direction worth exploring in the near future. This would mean going from *‘serotype predicts prevalence’* [Weinberger et al., 2009], to *‘serotype predicts interactions predicts prevalence’* paradigm. Similarly, exploring microbial ecosystem’s invasion fitness in pneumococcus on the basis of trait-based interactions could be a ground for verifying theoretical expectations from other more general ecological invasibility models [Hui et al., 2021]. Such bottom-up approaches would help also to establish whether the mode of serotype coexistence is the one predicted by our estimation results of Order 1 or Order 2, suggesting principles in favor of oscillatory vs. stable coexistence, which at this point remain uncertain.

### Future data and applications in other systems

We provide a framework based on coupled mechanistic invasion fitness networks, able to identify and quantify the common ecological underpinning of multiple coexistence patterns (linked via a resource gradient, here *µ*). We believe this is the first large-scale investigation of between-serotype interactions in pneumococcus, something that would be computationally prohibitive using the full SIS model with coinfection and multiple strains. Our spirit and results fall into a broad ecological topic increasingly interested on multi-species interactions and assembly in variable environments [Chesson, 1994, Grilli, 2020, Lewnard and Hanage, 2019]. Our estimation results on the rescaled interaction matrix are similar to the ones obtained by Tokita [2023], applied to murine microbiota time-series using the replicator system. We believe applications of our methodological framework to microbiota snapshots in different hosts, connected via a mechanistic gradient, as proposed in [Gjini and Madec, 2023], could be a next natural step for applications.

## Supporting information

Integrated Supplementary Material

Extended results files

## Acknowledgements

We thank Ermanda Dekaj who assisted with the raw data collection in the initial stages of this manuscript. This project was supported by the Portuguese Foundation for Science and Technology, (FCT) through grant 2022.03060.PTDC.

## Supplementary material

**S1** Data files. **S2** Code files. **S3 Text** Order 1, details of parameter estimation. **S4 Text** Order 2, details of parameter estimation. **S5** Extended results files. **S6** Supporting figures and tables. **S7** Estimation process with equal *µ* = 6 across sites.

## Notes

https://github.com/lthminhthao/ReplicatorParameters-Estimation/tree/main

### Competing Interest Statement

The authors have declared no competing interest.

### Summary of Updates

- Clarification of links with previous work - More details on the PSO algorithm - More interpretation on the estimated interactions - More details on the method limitations and future extensions

## References

T. Adebanjo, F. C. Lessa, H. Mucavele, B. Moiane, A. Chauque, F. Pimenta, S. Massora, M. d. G. Carvalho, C. G. Whitney, and B. Sigauque. Pneumococcal carriage and serotype distribution among children with and without pneumonia in Mozambique, 2014-2016. PloS one, 13(6):e0199363, 2018.

R. Aguas, I. Dorigatti, L. Coudeville, C. Luxemburger, and N. Ferguson. Cross-serotype interactions and disease outcome prediction of dengue infections in Vietnam. Scientific Reports, 9 (1):9395, 2019.

D. Akman, O. Akman, and E. Schaefer. Parameter estimation in ordinary differential equations modeling via particle swarm optimization. Journal of Applied Mathematics, 2018(1):9160793, 2018.

S. Alizon. Co-infection and super-infection models in evolutionary epidemiology. Interface focus, 3(6):20130031, 2013.

S. Alizon, J. C. de Roode, and Y. Michalakis. Multiple infections and the evolution of virulence. Ecology letters, 16(4):556–567, 2013.

S. Alizon, C. L. Murall, E. Saulnier, and M. T. Sofonea. Detecting within-host interactions from genotype combination prevalence data. Epidemics, 29:100349, 2019.

S. Allesina and J. M. Levine. A competitive network theory of species diversity. Proceedings of the National Academy of Sciences, 108(14):5638–5642, 2011.

S. Allesina and S. Tang. Stability criteria for complex ecosystems. Nature, 483(7388):205–208, 2012.

M. Alshawaqfeh, E. Serpedin, and A. B. Younes. Inferring microbial interaction networks from metagenomic data using SgLV-EKF algorithm. BMC genomics, 18:1–12, 2017.

J.-P. Aubin and K. Sigmund. Permanence and viability. Journal of Computational and Applied Mathematics, 22(2-3):203–209, 1988.

K. Auranen, J. Mehtälä, A. Tanskanen, and M. S. Kaltoft. Between-strain competition in acquisition and clearance of pneumococcal carriage? epidemiologic evidence from a longitudinal study of day-care children. American journal of epidemiology, 171(2):169–176, 2010.

T. Bedford, A. Rambaut, and M. Pascual. Canalization of the evolutionary trajectory of the human influenza virus. BMC biology, 10:1–12, 2012.

S. D. Bentley, D. M. Aanensen, A. Mavroidi, D. Saunders, E. Rabbinowitsch, M. Collins, K. Donohoe, D. Harris, L. Murphy, M. A. Quail, et al. Genetic analysis of the capsular biosynthetic locus from all 90 pneumococcal serotypes. PLoS genetics, 2(3):e31, 2006.

A. K. Chaturvedi, H. A. Katki, A. Hildesheim, A. C. Rodríguez, W. Quint, M. Schiffman, L.-J. Van Doorn, C. Porras, S. Wacholder, P. Gonzalez, et al. Human papillomavirus infection with multiple types: pattern of coinfection and risk of cervical disease. The Journal of infectious diseases, 203(7):910–920, 2011.

P. Chesson. Multispecies competition in variable environments. Theoretical population biology, 45(3):227–276, 1994.

S. Cobey, E. B. Baskerville, C. Colijn, W. Hanage, C. Fraser, and M. Lipsitch. Host population structure and treatment frequency maintain balancing selection on drug resistance. Journal of The Royal Society Interface, 14(133):20170295, 2017.

M. L. Cody and J. M. Diamond. Ecology and evolution of communities. Harvard University Press, 1975.

K. Z. Coyte, J. Schluter, and K. R. Foster. The ecology of the microbiome: networks, competition, and stability. Science, 350(6261):663–666, 2015.

E. Dekaj and E. Gjini. Pneumococcus and the stress-gradient hypothesis: A trade-off links R0 and susceptibility to co-colonization across countries. Theoretical Population Biology, 156:77–92, 2024.

R. Eberhart and J. Kennedy. Particle swarm optimization. In Proceedings of the IEEE international conference on neural networks, volume 4, pages 1942–1948. Citeseer, 1995.

M. Garcia Quesada, M. E. Peterson, J. C. Bennett, K. Hayford, S. L. Zeger, Y. Yang, M. K. Hetrich, D. R. Feikin, A. Von Gottberg, M. van der Linden, et al. Serotype distribution of remaining pneumococcal meningitis in the mature PCV10/13 period: Findings from the PSERENADE Project. Microorganisms, 9 (4), 2021.

K. A. Geno, G. L. Gilbert, J. Y. Song, I. C. Skovsted, K. P. Klugman, C. Jones, H. B. Konradsen, and M. H. Nahm. Pneumococcal capsules and their types: past, present, and future. Clinical microbiology reviews, 28(3):871–899, 2015.

E. Gjini. Geographic variation in pneumococcal vaccine efficacy estimated from dynamic modeling of epidemiological data post-PCV7. Scientific Reports, 7(1):1–16, 2017.

E. Gjini and S. Madec. The ratio of single to co-colonization is key to complexity in interacting systems with multiple strains. Ecology and Evolution, 11(13):8456–8474, 2021.

E. Gjini and S. Madec. Towards a mathematical understanding of invasion resistance in multispecies communities. Royal Society Open Science, 10(11):231034, 2023.

E. Gjini, C. Valente, R. Sa-Leao, and M. G. M. Gomes. How direct competition shapes coexistence and vaccine effects in multi-strain pathogen systems. Journal of Theoretical Biology, 388:50–60, 2016.

J. Gog and J. Swinton. A status-based approach to multiple strain dynamics. Journal of mathematical biology, 44(2):169–184, 2002.

J. Grilli. Macroecological laws describe variation and diversity in microbial communities. Nature communications, 11(1):1–11, 2020.

Z. B. Harboe, H.-C. Slotved, H. B. Konradsen, and M. S. Kaltoft. A pneumococcal carriage study in Danish pre-school children before the introduction of pneumococcal conjugate vaccination. The open microbiology journal, 6:40, 2012.

J. Hofbauer and K. Sigmund. Evolutionary games and population dynamics. Cambridge university press, 1998.

J. Hu, D. R. Amor, M. Barbier, G. Bunin, and J. Gore. Emergent phases of ecological diversity and dynamics mapped in microcosms. Science, 378(6615):85–89, 2022.

S. P. Hubbell. The unified neutral theory of biodiversity and biogeography (MPB-32). Princeton University Press, 2011.

C. Hui, D. M. Richardson, P. Landi, H. O. Minoarivelo, H. E. Roy, G. Latombe, X. Jing, P. J. CaraDonna, D. Gravel, B. Beckage, et al. Trait positions for elevated invasiveness in adaptive ecological networks. Biological Invasions, 23:1965–1985, 2021.

M. Imöhl, R. R. Reinert, and M. van der Linden. Regional differences in serotype distribution, pneumococcal vaccine coverage, and antimicrobial resistance of invasive pneumococcal disease among German federal states. International journal of medical microbiology, 300(4):237–247, 2010.

C. Jacquet, C. Moritz, L. Morissette, P. Legagneux, F. Massol, P. Archambault, and D. Gravel. No complexity–stability relationship in empirical ecosystems. Nature communications, 7(1): 12573, 2016.

R. Kandasamy, M. Gurung, A. Thapa, S. Ndimah, N. Adhikari, D. R. Murdoch, D. F. Kelly, D. E. Waldron, K. A. Gould, S. Thorson, et al. Multi-serotype pneumococcal nasopharyngeal carriage prevalence in vaccine naive Nepalese children, assessed using molecular serotyping. PloS one, 10(2):e0114286, 2015.

K. Koelle, S. Cobey, B. Grenfell, and M. Pascual. Epochal evolution shapes the phylodynamics of interpandemic influenza a (H3N2) in humans. Science, 314(5807):1898–1903, 2006.

T. M. T. Le and S. Madec. Spatiotemporal evolution of coinfection dynamics: a reaction– diffusion model. Journal of Dynamics and Differential Equations, 37(1):317–362, 2025.

T. M. T. Le, S. Madec, and E. Gjini. Disentangling how multiple traits drive 2 strain frequencies in SIS dynamics with coinfection. Journal of Theoretical Biology, 538:111041, 2022.

T. M. T. Le, E. Gjini, and S. Madec. Quasi-neutral dynamics in a coinfection system with N strains and asymmetries along multiple traits. Journal of Mathematical Biology, 87(3):48, 2023.

J. A. Lewnard and W. P. Hanage. Making sense of differences in pneumococcal serotype replacement. The Lancet Infectious Diseases, 19(6):e213–e220, 2019.

J. Lin, V. Andreasen, and S.A. Levin. Dynamics of influenza a drift: the linear three-strain model. Mathematical biosciences, 162(1-2):33–51, 1999.

M. Lipsitch, C. Colijn, T. Cohen, W. P. Hanage, and C. Fraser. No coexistence for free: neutral null models for multistrain pathogens. Epidemics, 1(1):2–13, 2009.

M. Lipsitch, O. Abdullahi, A. D’Amour, W. Xie, D. M. Weinberger, E. T. Tchetgen, and J. A. G. Scott. Estimating rates of carriage acquisition and clearance and competitive ability for pneumococcal serotypes in Kenya with a Markov transition model. Epidemiology (Cambridge, Mass.), 23(4):510, 2012.

C. Lo and R. Marculescu. Inferring microbial interactions from metagenomic time-series using prior biological knowledge. In Proceedings of the 8th ACM International Conference on Bioinformatics, Computational Biology, and Health Informatics, pages 168–177, 2017.

R. MacArthur. On the relative abundance of species. The American Naturalist, 94(874):25–36, 1960.

S. Madec and E. Gjini. Predicting n-strain coexistence from co-colonization interactions: epidemiology meets ecology and the replicator equation. Bulletin of Mathematical Biology, 82 (142), 2020.

I. Man, K. Auranen, J. Wallinga, and J. A. Bogaards. Capturing multiple-type interactions into practical predictors of type replacement following human papillomavirus vaccination. Philosophical Transactions of the Royal Society B, 374(1773):20180298, 2019.

I. Man, E. Benincá, M. E. Kretzschmar, and J. A. Bogaards. Reconstructing multi-strain pathogen interactions from cross-sectional survey data via statistical network inference. Journal of the Royal Society Interface, 20(205):20220912, 2023.

S. Marino, N. T. Baxter, G. B. Huffnagle, J. F. Petrosino, and P. D. Schloss. Mathematical modeling of primary succession of murine intestinal microbiota. Proceedings of the National Academy of Sciences, 111(1):439–444, 2014.

R. M. May. Will a large complex system be stable? Nature, 238(5364):413–414, 1972.

K. S. McCann. The diversity–stability debate. Nature, 405(6783):228–233, 2000.

N. Mejlhede, B. V. Pedersen, M. Frisch, and A. Fomsgaard. Multiple human papilloma virus types in cervical infections: competition or synergy? Apmis, 118(5):346–352, 2010.

M. R. Moore. Rethinking replacement and resistance, 2009.

I. Motomura. On the statistical treatment of communities. Zoological Magazine, Tokyo, 44: 379–383, 1932.

A. Raß, M. Schmitt, and R. Wanka. Explanation of stagnation at points that are not local optima in particle swarm optimization by potential analysis. In Proceedings of the Companion Publication of the 2015 Annual Conference on Genetic and Evolutionary Computation, GECCO Companion ‘15, page 1463–1464. Association for Computing Machinery, 2015.

H. Rodrigues, T. Pinto, R. Barros, L. Teixeira, and F. Neves. Pneumococcal nasopharyngeal carriage among children in Brazil prior to the introduction of the 10-valent conjugate vaccine: a culture-and pcr-based survey. Epidemiology & Infection, 145(8):1720–1726, 2017.

S. Shrestha, B. Foxman, D. M. Weinberger, C. Steiner, C. Viboud, and P. Rohani. Identifying the interaction between influenza and pneumococcal pneumonia using incidence data. Science translational medicine, 5(191):191ra84–191ra84, 2013.

R. R. Stein, V. Bucci, N. C. Toussaint, C. G. Buffie, G. Rätsch, E. G. Pamer, C. Sander, and J. B. Xavier. Ecological modeling from time-series inference: insight into dynamics and stability of intestinal microbiota. PLoS computational biology, 9(12):e1003388, 2013.

S. R. Tabatabaei, F. Fallah, F. Shiva, A. Shamshiri, M. Hajia, M. Navidinia, A. Karimi, and M. Rahbar. Multiplex pcr assay for detection of pneumococcal serotypes in nasopharyngeal samples of healthy children; Tehran, 2009-2010. Annual Research & Review in Biology, pages 3780–3790, 2014.

K. Tokita. Species abundance patterns in complex evolutionary dynamics. Physical review letters, 93(17):178102, 2004.

K. Tokita. Statistical mechanics of relative species abundance. Ecological Informatics, 1(3): 315–324, 2006.

K. Tokita. May’s dream: the interactions in biological networks were random matrices after all. In The Eighth International Symposium on BioComplexity, 2023.

M. van Baalen and M. W. Sabelis. The dynamics of multiple infection and the evolution of virulence. The American Naturalist, 146(6):881–910, 1995.

D. M. Weinberger, K. Trzciński, Y.-J. Lu, D. Bogaert, A. Brandes, J. Galagan, P. W. Anderson, R. Malley, and M. Lipsitch. Pneumococcal capsular polysaccharide structure predicts serotype prevalence. PLoS pathogens, 5(6):e1000476, 2009.

P. S. Wikramaratna, A. Kucharski, S. Gupta, V. Andreasen, A. R. McLean, and J. R. Gog. Five challenges in modelling interacting strain dynamics. Epidemics, 10:31–34, 2015.

Y. Xu, X. Zhou, W. Zheng, B. Cui, C. Xie, Y. Liu, X. Qin, and J. Liu. Serotype distribution, antibiotic resistance, multilocus sequence typing, and virulence factors of invasive and non-invasive streptococcus pneumoniae in Northeast China from 2000 to 2021. Medical Microbiology and Immunology, 213(1):12, 2024.

Y. Yoshino, T. Galla, and K. Tokita. Rank abundance relations in evolutionary dynamics of random replicators. Physical Review E—Statistical, Nonlinear, and Soft Matter Physics, 78 (3):031924, 2008.

