## Supplementary material for "Inference of pairwise interactions from strain frequency data across settings and context-dependent mutual invasibilities": Integrated Supplementary Material

#### S1 Data files

Frequencies and identities of serotypes in each study. We also provide the ordered list of the total 92 serotypes used to describe the global $\alpha_{ij}$ matrix. Please see uploaded dataset Github link.

#### S2 Code file

Nested PSO algorithm in Python notebook Github link. Please see uploaded notebooks .

#### S3 Text

##### Order 1: Details of estimation procedure (Brazil $\rightarrow$ Denmark $\rightarrow$ Mozambique $\rightarrow$ Iran $\rightarrow$ Nepal)

**Brazil ($n = 26, \mu = 9.85$).** The number of serotypes reported in Brazil is $n = 26$ and the ratio of single to co-colonization prevalence is $\mu = 9.85$. We start by estimating the matrix $26 \times 26$ $(\alpha_{ij})$ of Brazil. Since $\mu$ of Brazil is large, we estimate parameters such that the Brazil serotype frequencies correspond to the solution of the replicator dynamics at time $\tau = 50$ with the same frequencies for all initial values $z_i(0) = 1/n$, $1 \leq i \leq 26$. The target function for PSO algorithm satisfies that: i) For each matrix $(\alpha_{ij})$, we calculate the $\Lambda$ matrix of pairwise invasion fitness, which allows us to have the solution for replicator dynamics at $\tau = 50$. ii) For that solution, we sort it in descending order and compute the error between it and observation data of Brazil. We need this error to be minimal as possible. After fitting the parameters with PSO algorithm in range $[-10, 10]$ to attain the minimum error, we re-order the strains' name to have the $\alpha$ matrix with the right strains corresponding.

**Denmark ($n = 33, \mu = 10.23$)| *Brazil.*** The number of pneumococcus serotypes reported in Denmark is 33, and the ratio $I/D$ of single to co-colonization is $\mu = 10.23$. We need to estimate the matrix $33 \times 33$ $(\alpha_{ij})$ of Denmark. Since $\mu$ of Denmark is large, we estimate parameters such that the data observation (Denmark serotype abundances) is the solution of the replicator dynamics at time $\tau = 50$ with the same frequencies for all initial values $z_N(0)$, $1 \leq N \leq 33$. i) First, from $\alpha_{ij}$'s estimated from Brazil, we insert as fixed the $\alpha_{ij}$'s values already estimated for the serotype pairs common between the two sites. ii) Then we apply the PSO algorithm in the range $[-10, 10]$ to estimate the remaining 833 $\alpha_{ij}$'s such that the error between data observation

$z^*_{\text{Denmark}}$ and the solution of replicator dynamics (corresponding to $\Lambda$ matrix calculated from $(\alpha_{ij})$) at time $\tau = 50$ $\mathbf{z}(50)$ attains as low value as possible.

**Mozambique** ($n = 45, \mu = 16.8$) | ***Brazil, Denmark.*** The number of strains reported in Mozambique is 45; and the ratio of single to co-colonization is $\mu = 16.8$. We need to estimate the matrix $45 \times 45$ $(\alpha_{ij})$ of Mozambique. Since $\mu$ of Mozambique is large, we estimate parameters such that the data observation (Mozambique prevalence) is the solution of the replicator dynamics at time $\tau = 50$ with the same frequencies for all initial values $z_N(0)$, $1 \leq N \leq 45$, as what we did for Brazil and Denmark. The estimation process is as follows: i) From $\alpha_{ij}$'s estimated in Brazil and Denmark, in Mozambique's $\alpha_{ij}$ matrix, we insert 810 values of $\alpha_{ij}$'s already estimated in two previous processes. ii) Use PSO algorithm wtih range $[-10, 10]$ to estimate 1215 remaining $\alpha_{ij}$'s such that the error between data observation $z^*_{\text{Mozambique}}$ and the solution of replicator dynamics with the $\Lambda$ for Mozambique, at time $\tau = 50$, $\mathbf{z}(50)$ attains its minimum.

**Iran** ($n = 30, \mu = 0.93$) | ***Brazil, Denmark, Mozambique.*** The number of serotypes reported in Iran is 30; and $\mu = 0.93$. We need to estimate the matrix $30 \times 30$ $(\alpha_{ij})$ of Iran. Since $\mu$ of Iran is not large, we estimate parameters such that the data observation (Iran prevalence) is the stable equilibrium of the replicator dynamics. Our process are: i) From $\alpha_{ij}$'s estimated from previous epidemiological sites, in Iran's $\alpha_{ij}$ matrix, we insert 288 values of common $\alpha_{ij}$'s. ii) Then we calculate $\Lambda$ matrix and use PSO algorithm in range $[-10, 10]$ to estimate 612 remaining $\alpha_{ij}$'s such that data observation $z^*$ satisfies: $\left\|\Lambda z^* - {z^*}^T \Lambda z^*\right\|_2$ attains its minimum and the equilibrium is stable, i.e. the Jacobian matrix corresponding to the equilibrium $\Lambda z^*$ has all strictly negative real-part eigenvalues.

**Nepal** ($n = 69, \mu = 3.95$) | ***Brazil, Denmark, Mozambique, Iran.*** The number of strains in Nepal is $n = 69$; and $\mu = 3.95$. We need to estimate the matrix $69 \times 69$ $(\alpha_{ij})$ of Nepal. Since $\mu$ of Nepal is neither too large and nor too small, we estimate parameters such that the data observation (Nepal relative serotype prevalences) are i) the equilibrium of the replicator dynamics, or ii) a snapshot of the dynamics. The process is similar to the parameter estimation for Iran's $\alpha_{ij}$'s: i) From $\alpha_{ij}$'s estimated in Brazil, Denmark, Mozambique, and Iran, in Nepal's $\alpha_{ij}$ matrix, we insert 1901 values estimated before. ii) SS: Calculate $\Lambda$ matrix and use PSO algorithm in range $[-10, 10]$ to estimate 2860 remaining $\alpha_{ij}$'s such that data observation $z^*$ satisfies $\left\|\Lambda z^* - {z^*}^T \Lambda z^*\right\|_2$ attains its minimum and the equilibrium is stable, i.e. the Jacobian matrix corresponding to the equilibrium $\Lambda z^*$ has all strictly negative real-part eigenvalues. In PSO algorithm, we choose the range $[-10, 10]$ for these 2860 $\alpha_{ij}$'s. iii) Dyn: Calculate $\Lambda$ matrix and use PSO algorithm in range $[-10, 10]$ to estimate 2860 remaining $\alpha_{ij}$'s such that data observation is a snapshot of the dynamics.

**Computation time** The total time for the estimation process in this order is approximately 84 to 96 hours, which is relatively lengthy, despite the estimation of Brazil's parameters taking only 3 hours. This extended duration is primarily due to the sequential estimation of $\alpha_{ij}$ parameters in ascending order of strain number. As illustrated in the Venn diagram (Figure 2) and the block matrix visualization (Figure 3), the subsequent steps involve estimating an increasing number of $\alpha_{ij}$ parameters, albeit with limited degrees of freedom, resulting in longer estimation times. The computation of eigenvalues for the Jacobian matrices of Iran and, especially, Nepal (with a $69 \times 69$ matrix) is particularly time-consuming due to the large dimensionality of the matrices and the reduced degrees of freedom in these cases.

### S4 Text

#### Order 2: Details on parameter estimation (Nepal $\rightarrow$ Brazil $\rightarrow$ Denmark $\rightarrow$ Mozambique $\rightarrow$ Iran)

**Nepal** $(n = 69, \mu = 3.95)$**.** The number of strains in Nepal is $n = 69$; and the ratio of single to co-colonization prevalence is $\mu = 3.95$. We need to estimate the matrix $69 \times 69$ $(\alpha_{ij})$ of serotypes in Nepal, assuming Nepal relative serotype prevalences are the stable equilibrium of the replicator dynamics. In this assumption (SS), we recall how to calculate $\Lambda$ matrix $\Lambda = \left(\lambda_i^j\right)_{1\leq i,j\leq 69}$ from knowing $(\alpha_{ij})$ matrix and use PSO algorithm in range $[-10, 10]$ to estimate 4761 $\alpha_{ij}$'s such that data observation $z^*$ satisfies $\left\|\Lambda z^* - z^{*T}\Lambda z^*\right\|_2$ attains its minimum and the Jacobian matrix of $\Lambda z^*$ has all strictly negative real-part eigenvalue. In PSO algorithm, we choose the range $[-10, 10]$ for these 4761 $\alpha_{ij}$'s. We want the $(\alpha_{ij})$ matrix in Order 2 to be comparable to Nepal's $\alpha_{ij}$ matrix in Order 1 in some meaningful way. Therefore, we scale the values of these 4761 $\alpha_{ij}$ elements so that the standard deviation of their distribution matches the standard deviation of the distribution of the Nepal $(\alpha_{ij})$ matrix in Order 1. This scaling is achieved by multiplying all values by approximately 9.8337.

**Brazil** $(n = 26, \mu = 9.85)|$ ***Nepal .*** The number of serotypes reported in Brazil is $n = 26$ and $\mu = 9.85$. We start by estimating the matrix $26 \times 26$ $(\alpha_{ij})$ of Brazil. Since $\mu$ of Brazil is large, we estimate parameters such that the Brazil serotype frequencies correspond to the solution of the replicator dynamics at time $\tau = 50$ with the same frequencies for all initial values $z_i(0) = 1/n$, $1 \leq i \leq 26$. The process is: i) First, from $\alpha_{ij}$'s estimated from Nepal, we insert as fixed the $\alpha_{ij}$'s values already estimated for the serotype pairs common between the two sites. ii) Then we apply the PSO algorithm in the range $[-10, 10]$ to estimate the remaining 147 $\alpha_{ij}$'s such that the error between data observation $z^*_{\text{Brazil}}$ and the solution of replicator dynamics (corresponding to $\Lambda$ matrix calculated from $(\alpha_{ij})$) at time $\tau = 50$ $\mathbf{z}(50)$ attains as low value as possible.

**Denmark** $(n = 33, \mu = 10.23)|$ ***Nepal, Brazil.*** The number of pneumococcus serotypes reported in Denmark is 33, and the ratio $I/D$ of single to co-colonization is $\mu = 10.23$. We need to estimate the matrix $33 \times 33$ $(\alpha_{ij})$ of Denmark. Since $\mu$ of Denmark is large, we estimate parameters such that the data observation (Denmark serotype abundances) is the solution of the replicator dynamics at time $\tau = 50$ with the same frequencies for all initial values $z_N(0)$, $1 \leq N \leq 33$. i) First, from $\alpha_{ij}$'s estimated from Nepal and Brazil, we insert as fixed the $\alpha_{ij}$'s values already estimated in two previous processes. ii) Then we apply the PSO algorithm in the range $[-10, 10]$ to estimate the remaining 305 $\alpha_{ij}$'s such that the error between data observation $z^*_{\text{Denmark}}$ and the solution of replicator dynamics (corresponding to $\Lambda$ matrix calculated from $(\alpha_{ij})$) at time $\tau = 50$ $\mathbf{z}(50)$ attains as low value as possible.

**Mozambique** $(n = 45, \mu = 16.8)$ | ***Nepal, Brazil, Denmark.*** The number of strains reported in Mozambique is 45; and the ratio of single to co-colonization is $\mu = 16.8$. We need to estimate the matrix $45 \times 45$ $(\alpha_{ij})$ of Mozambique. Since $\mu$ of Mozambique is large, we estimate parameters such that the data observation (Mozambique prevalence) is the solution of the replicator dynamics at time $\tau = 50$ with the same frequencies for all initial values $z_N(0)$, $1 \leq N \leq 45$, as what we did for Brazil and Denmark. The estimation process is: i) From $\alpha_{ij}$'s estimated in Nepal, Brazil and Denmark, in Mozambique's $\alpha_{ij}$ matrix, we insert 1611 values of $\alpha_{ij}$'s already estimated in three previous processes. ii) Use PSO algorithm with range $[-10, 10]$

to estimate 414 remaining $\alpha_{ij}$'s such that the error between data observation $z^*_{\text{Mozambique}}$ and the solution of replicator dynamics with the $\Lambda$ for Mozambique, at time $\tau = 50$, $\mathbf{z}(50)$ attains its minimum.

**Iran** ($n = 30, \mu = 0.93$) | ***Nepal, Brazil, Denmark, Mozambique.*** The number of serotypes reported in Iran is 30; and $\mu = 0.93$. We need to estimate the matrix $30 \times 30$ $(\alpha_{ij})$ of Iran. Since $\mu$ of Iran is not large, we estimate parameters such that the data observation (Iran prevalence) is the stable equilibrium of the replicator dynamics. Our process are: i) From $\alpha_{ij}$'s estimated from previous epidemiological sites, in Iran's $\alpha_{ij}$ matrix, we insert 331 values of common $\alpha_{ij}$'s. ii) Then we calculate $\Lambda$ matrix and use PSO algorithm in range $[-10, 10]$ to estimate 569 remaining $\alpha_{ij}$'s such that data observation $z^*$ satisfies: $\left\|\Lambda z^* - {z^*}^T \Lambda z^*\right\|_2$ attains its minimum and the Jacobian matrix corresponding to the equilibrium $\Lambda z^*$ has all strictly negative real-part eigenvalues.

**Computation time** The total time for the estimation process in this order is approximately 30 to 36 hours. This duration is primarily due to the initial estimation of Nepal's $\alpha_{ij}$ parameters, which significantly advances the degrees of freedom and requires relatively little time (about 3 to 4 hours) to estimate the 4761 parameters. As illustrated in the Venn diagram 2 and Figure 3, in the subsequent steps, only a small number of additional $\alpha_{ij}$ parameters need to be estimated, resulting in shorter estimation times for Brazil, Denmark, Mozambique, and Iran.

### S5 Extended results files (S5results.zip)

• Estimated $\alpha$-matrix in each site (from Order 1 and Order 2). Notice we encode by '50' all entries that were not estimated with this dataset. The range of all estimated $\alpha_{ij}$ instead should be $[-10, 10]$. In each folder of Order 1 and 2, there are these files: Iran.txt, Nepal.txt, Brazil.txt, Denmark.txt and Mozambique.txt
• Estimated global $\alpha$-matrix (Order 1 and Order 2).
In each folder of Order 1 and 2, there are these files: big_alpha_1.txt and big_alpha_2.txt corresponding to global $\alpha$-matrix in Order 1 and Order 2, respectively.
• Estimated $\Lambda$ invasion fitness matrix in each site (from Order 1 and Order 2)
In each folder of Order 1 and 2, there are these files: Iran_fitness.txt, Nepal_fitness.txt, Brazil_fitness.txt, Denmark_fitness.txt and Mozambique_fitness.txt corresponding to $\Lambda = \left(\lambda_i^j\right)$- invasion fitness matrices in each site in Order 1 and Order 2, respectively.
• Estimated global $\Lambda$ matrix (same $\mu$ value)

### S6 Supporting figures and tables

This section provides additional figures and tables that complement the main findings of the study. These supplementary materials offer detailed visualizations and data that support the analysis, helping to clarify complex relationships and enhance the understanding of the results presented in the main manuscript.

| Fit order | Site | Nr. of Strains | $\mu$ | Equilibrium assumption | Test pattern (Target function of PSO) | Best fit (RMSE) |
|---|---|---|---|---|---|---|
| 1 | Brazil | 26 | 9.85 | Dyn | $\|\|\mathbf{z}(50) - \mathbf{z}_{\text{Brazil}}\|\|_{\mathbb{R}^{26}}$ | 0.003 |
| 2 | Denmark | 33 | 10.23 | Dyn | $\|\|\mathbf{z}(50) - \mathbf{z}_{\text{Denmark}}\|\|_{\mathbb{R}^{33}}$ | 0.0094 |
| 3 | Mozambique | 45 | 16.8 | Dyn | $\|\|\mathbf{z}(50) - \mathbf{z}_{\text{Mozambique}}\|\|_{\mathbb{R}^{45}}$ | 0.0065 |
| 4 | Iran | 30 | 0.93 | SS | $\|\|\mathbf{z}_{\text{Iran}} - \mathbf{z}_{\text{stable equilibrium}}\|\|_{\mathbb{R}^{30}}$ | 0.0027 |
| 5 | Nepal | 69 | 3.95 | SS | $\|\|\mathbf{z}_{\text{Nepal}} - \mathbf{z}_{\text{stable equilibrium}}\|\|_{\mathbb{R}^{69}}$ | 0.0012 |
| | | | | Dyn | $\|\|\mathbf{z}(50) - \mathbf{z}_{\text{Nepal}}\|\|_{\mathbb{R}^{69}}$ | 0.0047 |

Table S1: **Parameter estimation process with Order 1.** We summarize features of parameter estimation process of five countries Brazil, Iran, Denmark, Mozambique and Nepal in this table. The first column (ordered) presents the order of countries in the nested estimation process. The last column gives the root-mean-squared error (per serotype). The stabilities of the dynamics in Iran and Nepal-SS are guaranteed. For these countries, we can calculate the Jacobian matrices corresponding to these equilibria and their eigenvalues. The maximum real parts of the eigenvalues for Iran and Nepal-SS are $-0.0016$ and $-0.00034$, respectively.

| Fit order | Site | Nr. of Strains | $\mu$ | Equilibrium assumption | Test pattern (Target function of PSO) | Best fit (RMSE) |
|---|---|---|---|---|---|---|
| 1 | Nepal | 69 | 3.95 | SS | $\|\|\mathbf{z}_{\text{Nepal}} - \mathbf{z}_{\text{stable equilibrium}}\|\|_{\mathbb{R}^{69}}$ | 0.000019 |
| 2 | Brazil | 26 | 9.85 | Dyn | $\|\|\mathbf{z}(50) - \mathbf{z}_{\text{Brazil}}\|\|_{\mathbb{R}^{26}}$ | 0.0024 |
| 3 | Denmark | 33 | 10.23 | Dyn | $\|\|\mathbf{z}(50) - \mathbf{z}_{\text{Denmark}}\|\|_{\mathbb{R}^{33}}$ | 0.0046 |
| 4 | Mozambique | 45 | 16.8 | Dyn | $\|\|\mathbf{z}(50) - \mathbf{z}_{\text{Mozambique}}\|\|_{\mathbb{R}^{45}}$ | 0.0047 |
| 5 | Iran | 30 | 0.93 | SS | $\|\|\mathbf{z}_{\text{Iran}} - \mathbf{z}_{\text{stable equilibrium}}\|\|_{\mathbb{R}^{30}}$ | 0.0007 |

Table S2: **Parameter estimation process with Order 2.** We summarize features of parameter estimation process of five countries Brazil, Iran, Denmark, Mozambique and Nepal in this table. The first column (ordered) presents the order of countries in the nested estimation process. The last column gives the root-mean-squared error (per serotype). Similarly to the Order 1, in this Order 2, the stabilities of the dynamics in Nepal-Case 1 and Iran are guaranteed. For these countries, we can calculate the Jacobian matrices corresponding to these equilibria and their eigenvalues. The maximum real parts of the eigenvalues for Nepal-SS and Iran are $-0.024$ and $-0.051$, respectively.

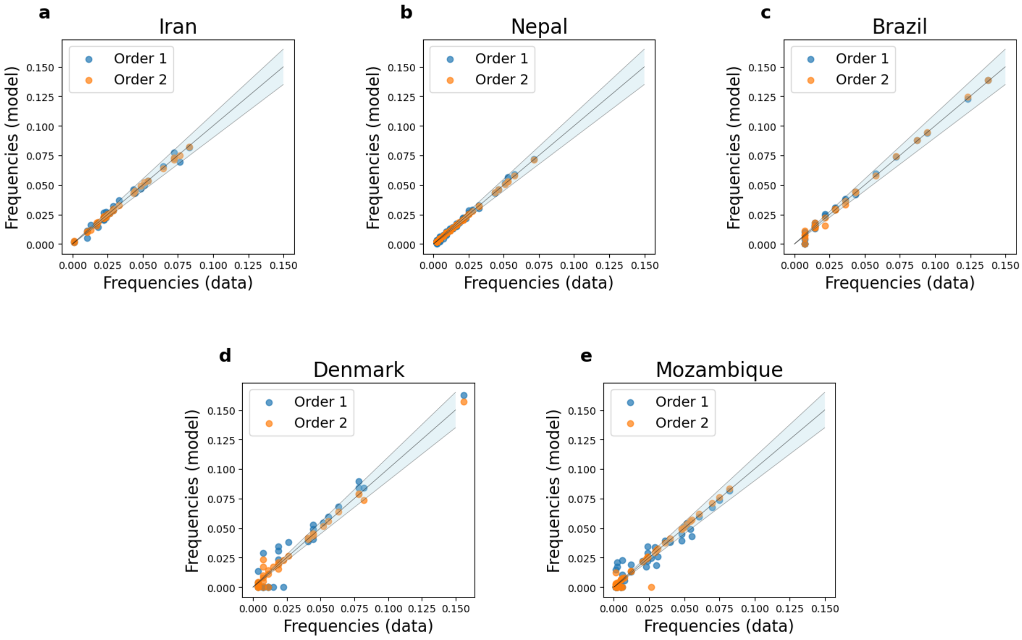


Figure S1: **Scatter plot of observed data versus predicted data in two approaches.** The blue and orange clouds in the five figures represent the scatter plots visualizing the estimation errors for predicted data for the five countries from Order 1 and Order 2, respectively. The mean square errors are shown in Tables S1 and S2 The shaded regions denote the acceptable regions corresponding to an estimation error of 10%, where the estimation is sufficiently accurate.

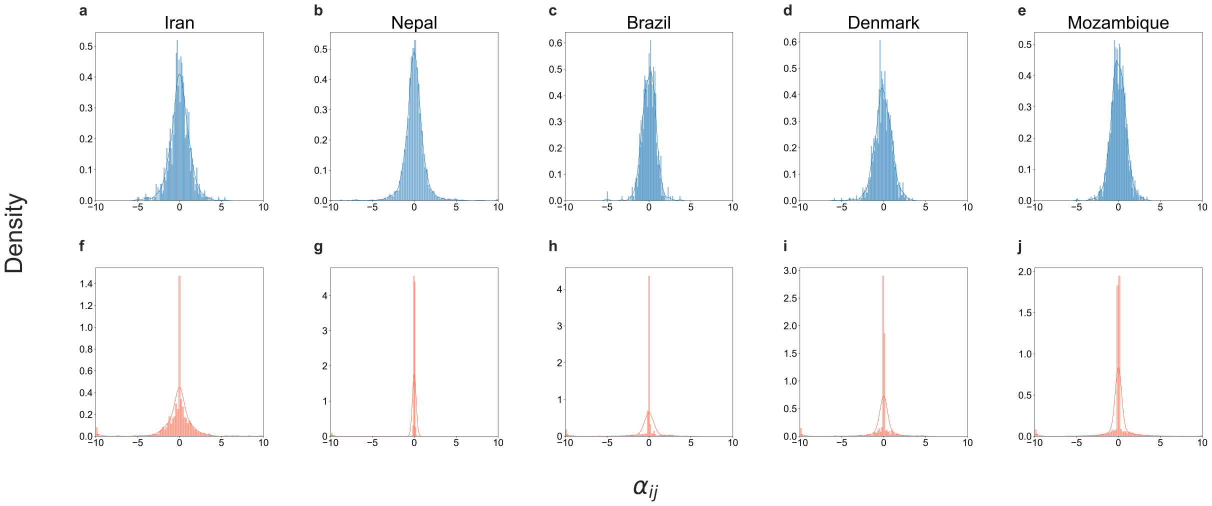


Figure S2: **Site-specific estimated distributions of the rescaled co-colonization interaction coefficients ($\alpha_{ij}$) in 5 epidemiological contexts of Streptococcus pneumoniae colonization for a total of 92 serotypes.** The upper panel (a-e) presents distributions of ($\alpha_{ij}$) for the countries Iran, Nepal, Brazil, Denmark, and Mozambique, obtained via Order 1 data fitting. These countries are arranged in ascending order of their single-to-coinfection ratio $\mu$. The bottom panel (f-j) illustrates distributions of the all $\alpha_{ij}$'s estimated via Order 2 data fitting. It is evident that, although in Order 2 we scaled the $\alpha_{ij}$ matrix of Nepal so that the standard deviation of the $\alpha_{ij}$ distribution matches that of Order 1, the distribution of $\alpha_{ij}$ in all five countries remains significantly more concentrated near zero compared to Order 1.

#### S7 Estimation process with uniform $\mu = 6$ across multiple countries (under the Stress-Gradient Hypothesis)

In this case, in line with the suggestion in [Dekaj and Gjini, 2024], we attempt parameter estimation under the assumption that $\mu = 6$ is consistent across all countries: Iran, Nepal, Brazil, Denmark, and Mozambique. Given that $\mu = 6$ is relatively large, we consider the data observations as snapshots of the dynamics at $\tau = 50$ for all countries. In this case, the pairwise invasion fitness coefficients are the same everywhere, only the strain composition changes.

**Site ordering for the nested $\alpha_{ij}$ estimation** Simultaneous estimation in this case should be slightly more advantageous than before, because under the SGH hypothesis, the invasion fitnesses $\lambda_i^j$ are equal across all countries allowing us to compute the corresponding $\Lambda$-matrix for this large $\alpha$-matrix at only one time . For each country, we extract the relevant portion of the fitness matrix (based on which serotype pairs are observed) to simulate the dynamics and calculate the error estimations. The objective function for this optimization process is the sum of all error estimations across five sites . However, this approach is still computationally intensive and requires significant computational resources as mentioned in the Discussion. Therefore, for practical application, we needed to adopt a sequential estimation approach also here: e.g. Order 1 or Order 2 (from before).

After some preliminary tests [1], in this section, we choose to estimate the $\alpha_{ij}$ parameters following again Order 1: the order Brazil $\rightarrow$ Denmark $\rightarrow$ Mozambique $\rightarrow$ Iran $\rightarrow$ Nepal, based on the increasing number of strains in subsequent countries.

A summary of these results is provided in Table S3 and in the Figures S3, S4 and S5.

| | Iran | Nepal | Brazil | Denmark | Mozambique |
|---|---|---|---|---|---|
| Best fit (RMSE) | 0.0084 | 0.0076 | 0.0036 | 0.0062 | 0.0082 |
| % $z$ predictions within 10% error margin | 40% | 7.25% | 69.23% | 27.27% | 33.33% |
| Symmetry level of $(\alpha_{ij})$-matrix | -0.049 | 0.012 | 0.042 | 0.006 | 0.011 |
| Symmetry level of $\left(\lambda_i^j\right)$-matrix | -0.98 | -0.98 | -0.98 | -0.98 | -0.98 |

Table S3: **Summary of parameter estimation assuming same $\mu = 6$ everywhere and conserved $\lambda_i^j$ across countries (SGH).** The consistent $\mu$ value also explains the much larger root mean square error compared to the previous $\mu$-dependent estimation (Orders 1 and 2), and the estimation process takes 5 to 6 times longer than in Order 1 (only Order 1 is compared because it follows the same country estimation order). The near-zero symmetry level of the $(\alpha_{ij})$ matrices indicates that these matrices are almost entirely random, and even more random than those estimated in Orders 1 and 2, see Table 2. Consequently, the corresponding $\Lambda$ invasion fitness matrices are nearly anti-symmetric, due to the consistently large value of $\mu = 6$.

**Fit and quality of fit** According to two first line of Table S3, under stress gradient hypothesis our algorithm does not fit well as in Order 1 and Order 2, however, it is able to fit certainly well the observed serotype prevalences in each study site (Figure S3), mean error being less than

[1] We conducted a heuristic experiment while estimating parameters under the stress gradient hypothesis, following the same country order as in Order 2. Notably, by the third step, it became nearly impossible to achieve parameter estimates with an error margin below 1.5%. This increased difficulty in parameter estimation for later countries made us decide for site ordering via Order 1.

1% for each ordering of data. We recall the reason is that, $\mu = 6$ everywhere, leading to the same invasion fitness $\lambda_i^j$ for identical pairs $(i, j)$ across all countries, which increases difficulty in parameter estimation for later countries, due to differing frequencies of the same strains across the five countries. Here we still use the same test pattern as in Order 1 and Order 2.

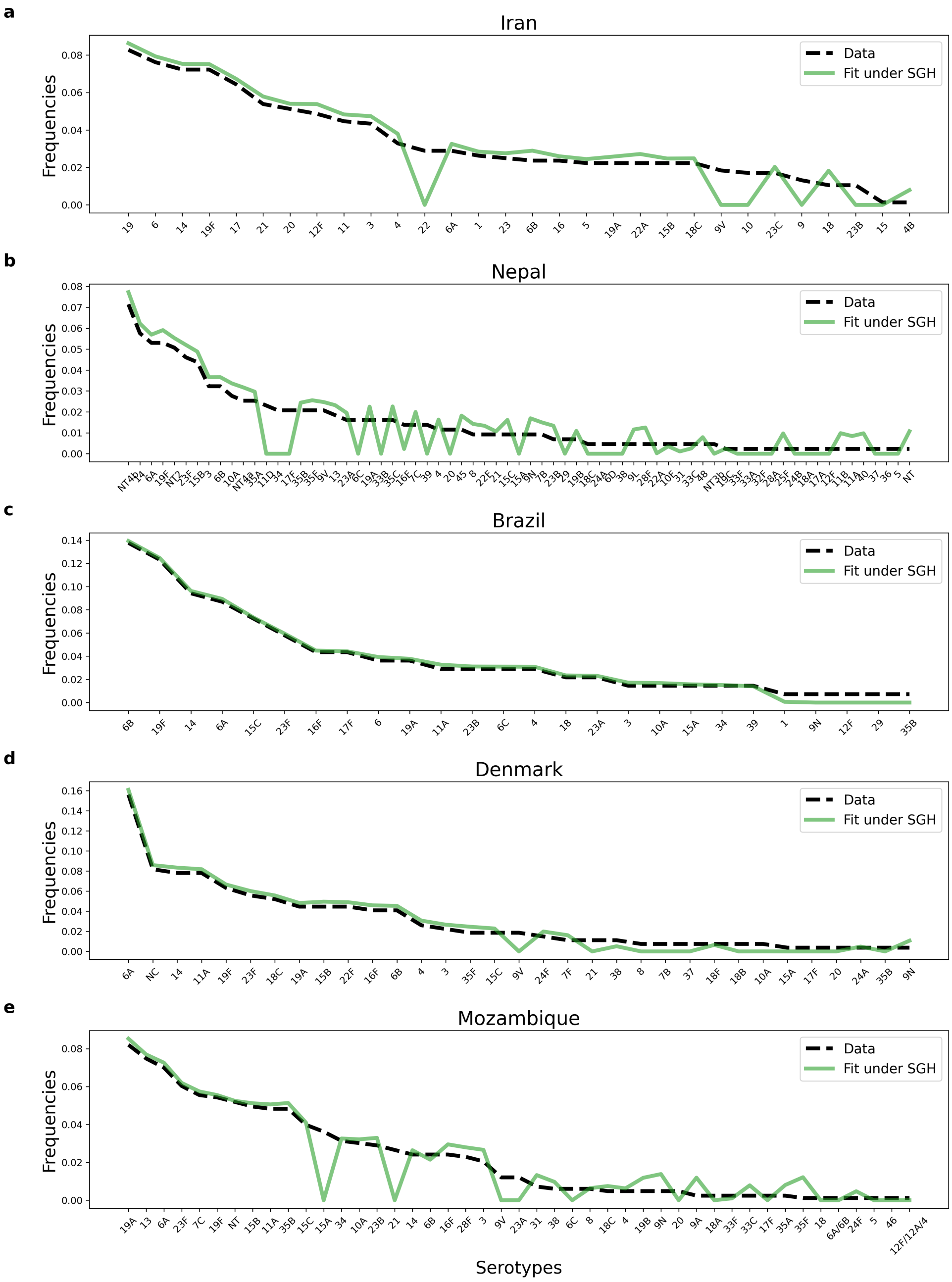


Figure S3: **Relative abundance data and model fits, assuming $\mu = 6$, for pneumococcus serotype frequencies in 5 epidemiological contexts.** We plot the data extracted from Denmark Harboe et al. [2012], Iran Tabatabaei et al. [2014], Brazil Rodrigues et al. [2017], Nepal Kandasamy et al. [2015], and Mozambique Adebanjo et al. [2018], as well as the model-predicted relative abundances (estimated from nested data fitting under stress-gradident hypothesis), ranked from the most prevalent to the least prevalent serotype.

**The mutual invasion fitness network that emerges** The pairwise invasion fitness matrix, which represents higher-order fitness influenced by contextual parameters such as the ratio of single to co-colonization, exhibits significant anti-symmetry for pneumococcal serotypes (characterized by predominantly $+-$ edges) under the stress gradient hypothesis and the chosen estimation method. This observation is consistent with the findings from Order 1 and is represented in Figure S4 and Figure S5. However, with $\mu = 6$ being lower than the $\mu$ values for Brazil, Denmark, and Mozambique in Order 1, we observe very few increased coexistence and bistability in this scenario. It is important to note that, with $\mu = 6$ applied universally, the pairwise fitnesses are identical for the same strain pairs across all countries. Consequently, plotting the network of common strains for one site effectively represents the network graph for all sites.

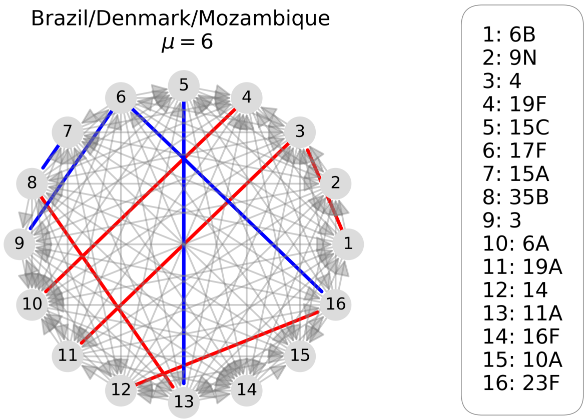


Figure S4: **Emergent invasion fitness network among 16 pneumococcus serotypes in different epidemiological settings, from parameter estimation under constant $\mu = 6$ (SGH, $\Lambda$ not context-dependent).** We show here the obtained mutual invasion fitness network for the co-occurring serotypes in Brazil, Denmark and Mozambique (only common serotypes), dominated by $+-$ edges (gray arrows) with rare coexistence (red) or bistability (blue). Here the pairwise outcomes are conserved across sites, only serotype composition changes.

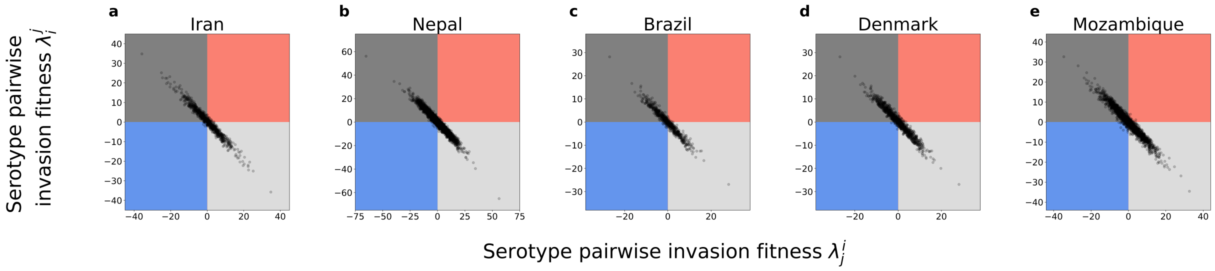


Figure S5: **Mutual invasion fitnesses between different pneumococcus serotype pairs across 5 countries resulting from estimation under $\mu = 6$ everywhere (the Stress-Gradient Hypothesis).** We plot the resulting pairwise invasion fitnesses between all serotypes present in each study site. For more details, we compute the percentages of pairwise outcomes (edges of the network), including exclusion of either strain, bistable outcome, and coexistence, for each country and each estimation order. We report the sequence triple (exclusion of either strain, bistability, coexistence) as the percentage among all pairwise outcomes: Iran $(89.8\%, 1.1\%, 5.8\%)$, Nepal $(92.3\%, 3.1\%, 3.2\%)$, Brazil $(89.9\%, 3.8\%, 2.4\%)$, Denmark $(89.3\%, 2.4\%, 5.3\%)$, Mozambique $(90.9\%, 3.4\%, 3.6\%)$. This confirms the symmetry level computed in Table S3.

**Comparison with Order 1** The order of countries estimated under the stress-gradient hypothesis follows the same sequence as in Order 1. Thus, it is valuable to compare the distributions of the $92 \times 92$ $\alpha$-matrices, as well as the individual $\alpha$-matrices for each country, across the two estimation approaches. Notably in Figure S6 and Figure S7, the distributions of the $92 \times 92$, $\alpha$-matrices in both Order 1 and the stress-gradient hypothesis (SGH) case are quite similar. Furthermore, for each site, the distributions of the corresponding $\alpha$-matrices show little deviation between the two estimation orders.

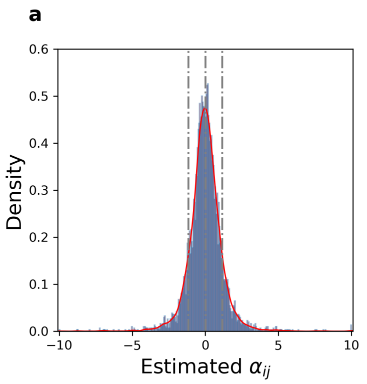


Figure S6: **Final global matrix distribution of rescaled interaction coefficients ($\alpha_{ij}$) for observed serotype pairs across 5 epidemiological contexts of Streptococcus pneumoniae colonization, after the nested fitting of cross-sectional data in Order 1 and under assuming SGH ($\mu = 6$ everywhere).** The figures illustrate distributions of the all $\alpha_{ij}$'s (73% of the entire matrix A) estimated in the two estimation approaches. We note that both distributions are very similar, appear bell-shaped, although the second case is a little bit more highly peaked at 0. The statistics for the two distributions are as follows. **(a)**- Order 1: Mean: 0.0038, Standard Deviation: 1.1610, Skewness: 0.1862, and Kurtosis: 13.2259. **(b)**- Under stress gradient hypothesis: Mean: 0.003, Standard Deviation: 0.8266, Skewness: $-0.4288$, and Kurtosis: 7.5355. For site-specific distributions see Figure S7.

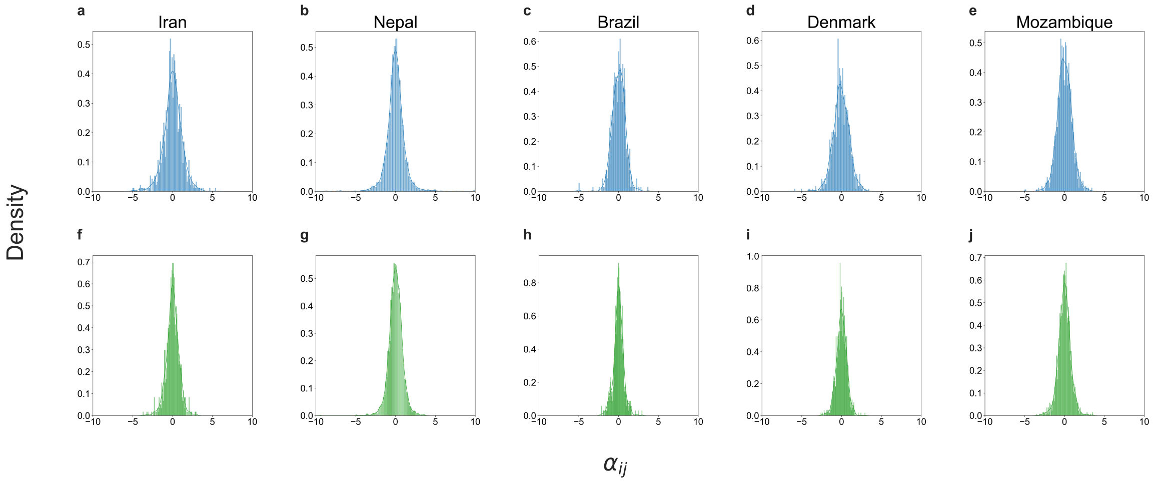


Figure S7: **Comparison of site-specific estimated distributions of the rescaled co-colonization interaction coefficients ($\alpha_{ij}$) in 5 epidemiological contexts of Streptococcus pneumoniae colonization for a total of 92 serotypes.** The upper panel (a-e) presents distributions of ($\alpha_{ij}$) for the countries Iran, Nepal, Brazil, Denmark, and Mozambique, obtained via Order 1 data fitting. These countries are arranged in ascending order of their single-to-coinfection ratio $\mu$. The bottom panel (f-j) illustrates distributions of the all $\alpha_{ij}$'s estimated via data fitting under stress gradient hypothesis.
